# Using language models and ontology topology to perform semantic mapping of traits between biomedical datasets

**DOI:** 10.1101/2022.08.02.502449

**Authors:** Yi Liu, Benjamin L Elsworth, Tom R Gaunt

## Abstract

**Motivation:** Human traits are typically represented in both the biomedical literature and large population studies as descriptive text strings. Whilst a number of ontologies exist, none of these perfectly represent the entire human phenome and exposome. Mapping trait names across large datasets is therefore time-consuming and challenging. Recent developments in language modelling have created new methods for semantic representation of words and phrases, and these methods offer new opportunities to map human trait names in the form of words and short phrases, both to ontologies and to each other. Here we present a comparison between a range of established and more recent language modelling approaches for the task of mapping trait names from UK Biobank to the Experimental Factor Ontology (EFO), and also explore how they compare to each other in direct trait-to-trait mapping.

**Results:** In our analyses of 1191 traits from UK Biobank with manual EFO mappings, the BioSentVec model performed best at predicting these, matching 40.3% of the manual mappings correctly. The BlueBERT-EFO model (finetuned on EFO) performed nearly as well (38.8% of traits matching the manual mapping). In contrast, Levenshtein edit distance only mapped 22% of traits correctly. Pairwise mapping of traits to each other demonstrated that many of the models can accurately group similar traits based on their semantic similarity.

**Availability and Implementation:** Our code is available at https://github.com/MRCIEU/vectology.

## Introduction

Population health and medical research are increasingly reliant on large population studies such as UK Biobank^1^, The Million Women Study^2^, Our Future Health^3^, The Million Veterans Program^4^, China Kadoorie Biobank^5^ and others to discover new predictive biomarkers and interventions. Such studies measure many thousands of phenotypic variables. Systematic analyses such as phenome-wide association studies (PheWAS)^6–8^ can describe relationships between thousands of variables, producing large datasets. However, many variables are inconsistently named across studies, and can prove difficult to map to each other or an existing ontology such as the Experimental Factor Ontology (EFO)^9^, Human Phenotype Ontology (HPO)^10^ or the Disease Ontology^11^. In parallel, the biomedical literature contains a wealth of data on human diseases, traits and risk factors described using free text (with some mappings to Medical Subject Headings; MeSH). Systematically integrating knowledge across these different datasets and domains would enable us to triangulate the evidence^12^ for different risk factor/disease combinations, but at the moment this is hindered by the inconsistencies in trait nomenclature.

The complexity of variable names is illustrated by UK Biobank, an internationally important population study that has collected a wealth of data on half a million people^1^. Examples of text labels for variables in UK Biobank include easily recognizable traits such as “systolic blood pressure” and disease names such as “coronary heart disease”. However, the study also includes more complex variables, including those derived from questionnaire data, including “able to walk or cycle unaided for 10 minutes” and “cough on most days”. An array of other variables also exist, including International Statistical Classification of Diseases and Related Health Problems 10th Revision (ICD10) codes such as “anaemia due to enzyme disorders” (D55) and “syncope and collapse” (R55), the former mapping directly to the EFO (EFO:0009529), but the latter not. Direct mapping by text matching to ontology terms is therefore not realistic, and whilst manual mapping to ontologies is sometimes appropriate, this is time consuming, especially if mapping to multiple different ontologies (which cover different domains of the human phenome and exposome).

Given this, there are four potential solutions to link two datasets based on their lists of trait (variable) names:

1. Manual mapping to an ontology to find shared terms between datasets
2. Using automated tools to map each variable to an ontology to find shared terms between datasets
3. Direct mapping of variables using a generalisable text embedding model to identify semantically similar terms
4. Direct mapping of variables using a bespoke model trained on the particular datasets to identify semantically similar terms

Each of these options has different strengths and weaknesses. Option 1 can only really be used in cases where the numbers of variables is low, or the requirement of human assigned ontological terms is essential. Option 2 relies on existing tools, such as OnToma^13^, Zooma^14^ or MetaMap Lite^15^ for common ontologies such as EFO^9^ and UMLS^16^. These rule-based tools can work well, but the mapping to ontology may identify a more generic parent term in the ontology losing valuable information in the process. Options 3 and 4 may offer benefits in mapping variables between datasets by avoiding the intermediate step of an ontology term (**Figure 1**).

**Figure 1:**
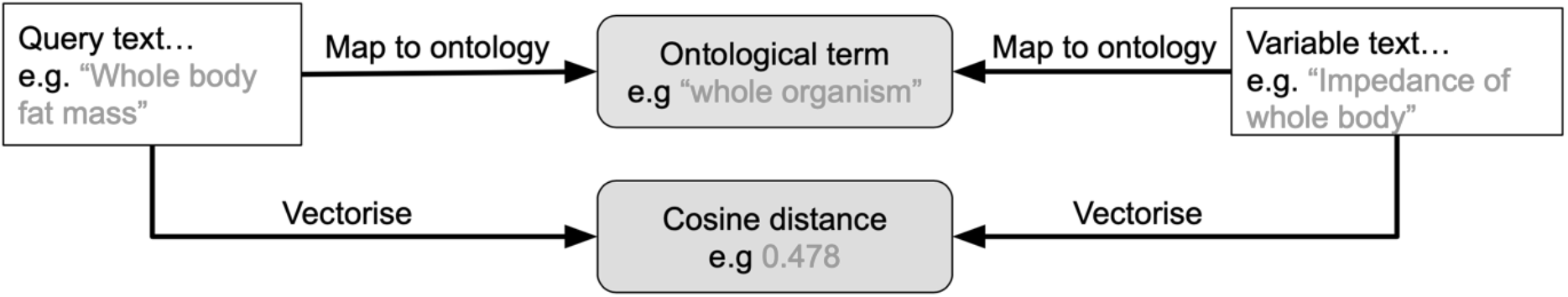
Example of potential benefits of using text embeddings to connect two biomedical strings compared to using a shared ontology

The development of methods based on text embeddings such as Word2Vec^17^, Sent2Vec^18^ and Doc2Vec^19^ have opened up the potential to map terms based on semantic similarity. These methods have been applied to data from the biomedical domain e.g. BioWordVec^20^ and BioSentVec^21^ and have been applied to real world problems^22–27^.

Further development to shallow / non-contextual text embeddings gives rise to contextualized methods such as the transformer model architecture^28^ and its implementation in language modelling (e.g. BERT^29^) applied in a range of contexts (e.g. GLUE^30^, BLUE^31^ and BLURB^32^). These models can be finetuned to tackle specific problems with great effect^33–37^. However, despite their merits, transformer models are slower and more resource intensive compared to the Word2Vec architecture.

Here we apply a range of text embedding methods and BERT language models (including one trained on EFO) to the problem of mapping biomedical variables (from UK Biobank) to an ontology (EFO) and compare their performance, strengths and weaknesses. We also illustrate the use of these models on a direct trait-to-trait mapping problem.

## System and Methods

### The EFO dataset

The **Experimental Factor Ontology (EFO)**^9^ contains parts of several biological ontologies as well as variables from many large scale databases. Whilst many other ontologies exist, this particular ontology is widely used for human traits and is well documented, so was considered a good choice for this evaluation. A version of the EFO data set was downloaded from the EBI RDF platform^37^ in March 2021 containing 25,390 terms. This was used for all subsequent analysis and is available in the supplementary material (**supplementary files S1 and S2**).

#### Ontology distance metric

To understand the relative distance between any two EFO terms and enable us to measure how well a trait was mapped we used the nxontology Python library^38^. By creating a parent child network of EFO terms we could compute a similarity measure between any pair of EFO terms and use this to create a measure of how close two terms are within the EFO hierarchy. For this analysis we used the Batet (parameter “*batet”* in the library) measure^39^ as this was developed using biomedical taxonomy data and produced good correlation results to manual biomedical concept comparisons. The measure ([0, 1]) is a ratio calculated from the shared and non-shared information between a pair of concepts, where the lower the score the less shared ancestry between the two ontology concepts have. From here on we will refer to this metric as the **EFO-Batet score**.

To create a nxontology instance, we provided the parent/child EFO edge data to the *NXOntology* class (**supplementary code block S1**).

#### Trait-to-trait mapping distance score

The different models use different approaches for measuring distance between text terms (**supplementary table S2**). For simplicity we refer to these metrics (edit distance, cosine similarity, semantic distance) as “trait similarity score” throughout.

### Mapping methods

We used a range of existing string comparison language models representing different approaches to language representation and different pre-training datasets to enable us to evaluate the impact of these differences on mapping performance.

#### String comparison methods

**Levenshtein edit distance ratio**^40^ was used to understand how well a basic string comparison performs. Using the *ratio()* function we obtained a measure of similarity between two strings.

**Zooma**^41^ is an established tool to map text to ontologies using a combination of curated mapping to existing data sets and standard text matching (the exact method is undocumented). For this analysis we utilised the Zooma API setting the “required” parameter set to “None” and “ontologies” parameter set to “efo” (**supplementary code block S2**) to avoid circularity.

#### Text embedding methods

**BioSentVec** is an established model created using sent2vec^18^, pre-trained on over 28 million titles and abstracts from PubMed^42^ and 2 million clinical notes from MIMIC III^43^. The BioSentVec^21^ model was downloaded from the project GitHub repository (https://github.com/ncbi-nlp/BioSentVec) and installed following the examples in the tutorial (https://github.com/ncbi-nlp/BioSentVec/blob/master/BioSentVec_tutorial.ipynb) (**supplementary code block 3**).

**Google Universal Sentence Encoder v4 (GUSE)** is a generalised text embedding model trained and optimised for sentence level tasks^44^. The model was trained on Wikipedia and other generalised texts with no focus on biomedical information. The model was downloaded from the project home page (https://tfhub.dev/google/universal-sentence-encoder/4) and implemented as described in the documented example (**supplementary code block S4**).

**spaCy** is a natural language processing platform which provides various tools, methods and pipelines, one of which is word embeddings^45^. The *en_core_web_lg* model was downloaded and installed as described in the documentation (https://spacy.io/usage/linguistic-features#vectors-similarity) (**supplementary code block S5**).

**ScispaCy** is built on spaCy and provides models for processing biomedical, scientific or clinical text^46^. The en_*core*_sci_lg model was downloaded and installed as described in the documentation (https://allenai.github.io/scispacy). The model is accessed via the same spaCy methods as above.

**BlueBERT**^31^ (NCBI_BERT_pubmed_mimic_uncased_L-12_H-768_A-12) and **BioBERT**^47^ (biobert_v1.1_pubmed) are biomedical language model implementations based on the original BERT pretrained weights, with further language model training with biomedical corpora to improve language understanding tasks in the biomedical domain. For transformer models, the vector representation of the entity is computed as the average of the hidden state tensor of the *N* − 1 layer as a fixed representation of the tokenised sequence (i.e. the default strategy in^48^). These models were obtained from their respective model repositories, then served via the bert-as-service^48^ API (see **supplementary code block S6** for example usage and code repository for detailed set up).

#### Bespoke ontology classifier

In addition to established language models we also explored the effect of tailoring a transformer model to the EFO using transfer learning.

**BlueBERT-EFO** was developed by finetuning BlueBERT with an ontology entity alignment training process designed as a sequence classification task (for details see **supplementary text S1**). To create a similarity matrix of the entities, for each pair of terms the model produces a score representing the inferred ontology distance, where the lower number of steps between two entities as predicted by the model, the closer they are represented in an ontology graph. The model can be used for inference using the Huggingface Transformers^49^ package (see **supplementary code block S7** for example usage and code repository for detailed set up). **Supplementary table S2** shows a summary of the models and methods.

### Mapping to ontology (EFO)

To assess how the models perform when mapping biomedical variables to an existing ontology, we utilised the **EBI UK Biobank EFO dataset**^50^. This is a list of around 1,500 UK Biobank variables that have been manually mapped to EFO terms. In addition, each mapping has been assigned a mapping type (Exact, Broad, Narrow, Other). The original data set was modified in the following ways: first, any query that had been assigned multiple EFO terms was dropped. Second, exact matches were excluded as uninformative (i.e. the query term is identical to the EFO label). Third, due to our use of an EFO hierarchy distance method (EFO-Batet) we only included those rows containing an EFO term present in our parent/child EFO data set. Fourth, all EFO and variable terms were lower-cased. Lastly, duplicates were removed. These filtering steps created a data set with 1,191 entries (**supplementary file S3**). Supplementary table S1 displays the numbers of each by mapping type and a brief description of each mapping type as described in the original data set.

Using this dataset, we applied the models described above to conduct pairwise comparison between the UK Biobank variables and the EFO terms to measure their semantic similarity and ontology distance. Specifically, a UK Biobank variable *A* is associated with a manually mapped EFO term *a* in the source dataset, then for an EFO term *b*, we calculated the similarity score between *A* and *b* as well as the EFO-Batet distance score between *a* and *b*. Therefore for the variable of interest *A*, the results dataset gives us a measure of how close the top ranking (by a specific similarity score metric) EFO term predictions *b*_0_ … *b*_*N*_ are to the variable’s equivalent EFO representation *a* in the ontology space (by the EFO-Batet score).

### Direct trait-to-trait mapping

In some scenarios mapping trait names between two datasets directly (without using an ontology) might be preferable. To compare how the different methods perform when predicting the similarity between two biomedical variables we again used the EBI UK Biobank EFO dataset^50^. This time, we limited the entries to those labelled as “Exact” on the assumption that these would provide a better dataset for assessing pairwise distances, both semantically and using the same ontology based method^38^. Additional filtering steps were taken to create a dataset with one query per predicted EFO term, resulting in 530 entries (**supplementary file S4**). For the purposes of visualisation, we then manually selected a subset of 43 traits that represented a broad spectrum of variables, covering measurements, questionnaire data and disease (**supplementary file S5**). For each of the pre-trained models, pairwise cosine distances were generated for each query text. For Levenshtein, the similarity ratio was calculated as before. For BlueBERT-EFO, we generate the inferred ontology distance for each pair of terms. Whilst we were not mapping trait terms to an ontology, we also compared how close these pairs of traits are in the EFO for comparison using the EFO-Batet score for each pair of terms.

## Implementation

### Comparison to other approaches for automated mapping to ontology

#### Top ranking results

We first explored how well the top prediction of each method compared to the manual annotation (**Figure 2**). For results that exactly agree with the manual annotation (**Figure 2A**), the best performing methods were BioSentVec (40.3%), BlueBERT-EFO (38.8%), Zooma (37.2%) and ScispaCy (36.5%), the results of which were notably higher than those of the methods included in the analysis. Pairwise proportions Z-test results (supplementary Table S4) between each of the mapped proportions confirm that there is a notable difference between results of the best performing group and the those of the other methods, but the differences are minimal within the group (largest difference is between BioSentVec and ScispaCy, P-Value = 0.058).

**Figure 2:**
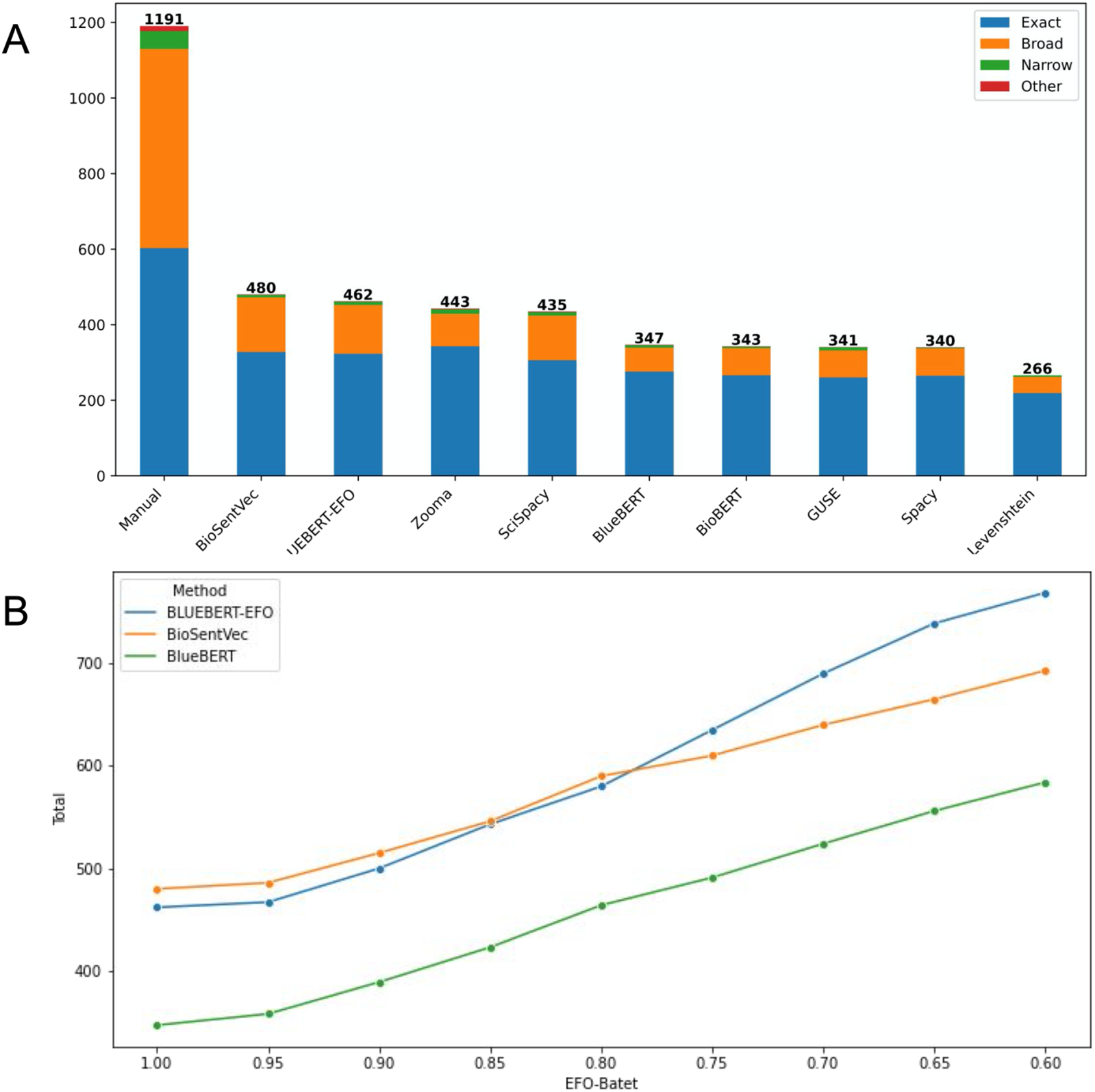
Distribution of top matching predictions. (A) Number of top matching predictions by MAPPING_TYPE. The Total bar contains all manual mappings, subdivided into Exact, Broad (parent term), Narrow (child term) and Other. Each other bar represents the number of traits exactly matched by the named method to the manual mapping for that trait, with the same subdivisions. (B) Total number of top matching predictions that are equal or above an EFO-Batet threshold, i.e. if a method produces greater number of matched predictions with a threshold closer to 1, greater number of predictions exhibit close ontology relationship to the manual mapping results. Points at EFO-Batet thresholds 1.0, 0.9, 0.8, 0.7 are equivalent to the Total values for each method in **Figures 2A, S1-S3**. Full results for all methods can be found in **Supplementary Figure S9**.

Whilst none of the methods exceeded 40.3% exact mapping, it is important to consider three key points: (1) some of the manual predictions are likely to be incorrect; (2) the methods and models used here to automate this approach are quick and easy to use, and would scale to a task size that would be impractical for manual annotation; (3) even the most sophisticated natural language processing models will struggle to predict the same result as a human, particularly in cases where the query string contains two un-linked entities, or even a negated term, e.g. “enduring personality changes not attributable to brain damage and disease”.

In some situations (e.g. a recommender of similar concepts), an exact match may not be required, and if the top prediction from a model is sufficiently close to the manual annotation, this may be a suitable result. We then examine how well the top predictions from a method align with the manual annotation in terms of their EFO-Batet score distance to the manual EFO terms. **Figure 2B** shows the aggregate results for the subset (see supplementary Figure S9 for full results) of methods over different range of EFO-Batet score threshold for top predictions to be included, from total number of top predictions that are strictly identical to manual annotation (threshold == 1, i.e. **Figure 2B**), to those that are sufficiently close to the manual annotation in the ontology space (e.g. threshold >= 0.9), and then to results with a greater ontology distance tolerance (e.g. threshold >= 0.6).

**Supplementary Figures S1-S3** show the detailed distributions for thresholds of 0.9, 0.8, and 0.7 respectively.

For inexact mapping results, BlueBERT-EFO and BioSentVec retrieved similar number of concepts that are close (e.g. under an EFO-Batet threshold of 0.9 or 0.8) to manual annotation, where notably greater number of predictions by BlueBERT-EFO have more ontology similarity to their manual annotation counterparts then the rest of the methods. In other words, BlueBERT-EFO as a finetuned model on BlueBERT with EFO structural information, is able to enhance the performance of the foundational BlueBERT to be on par with BioSentVec, and able to incorporate EFO knowledge on candidate retrieval.

#### Overall results for top N predictions

With methods that produce a distance or score, there may still be significant value in a set of top predictions (which we would expect to be enriched for related terms, and potentially contain the correct mapping term). We then investigated the distribution of EFO-Batet scores for both the top prediction (**Figure 3A**) and the top 10 predictions (weighted average EFO-Batet scores, **Figure 3B**), and the aggregate results of generalised top ranges, to determine which models prioritize the most relevant set of traits. As shown in **Figure 3A**, for top predictions BlueBERT-EFO is able to retrieve higher number of candidates that have high ontology relevance to the manual annotation (greater mass in the upper tail) and lower number of candidates that have low relevance (lower mass in the lower tail), which is also confirmed by the pairwise Kolmogorov-Smirnov two sample tests (supplementary table S4) on the statistical difference of its distribution to those of other methods (*P* − *Value* ≤ 3.3*E* − 09).

**Figure 3:**
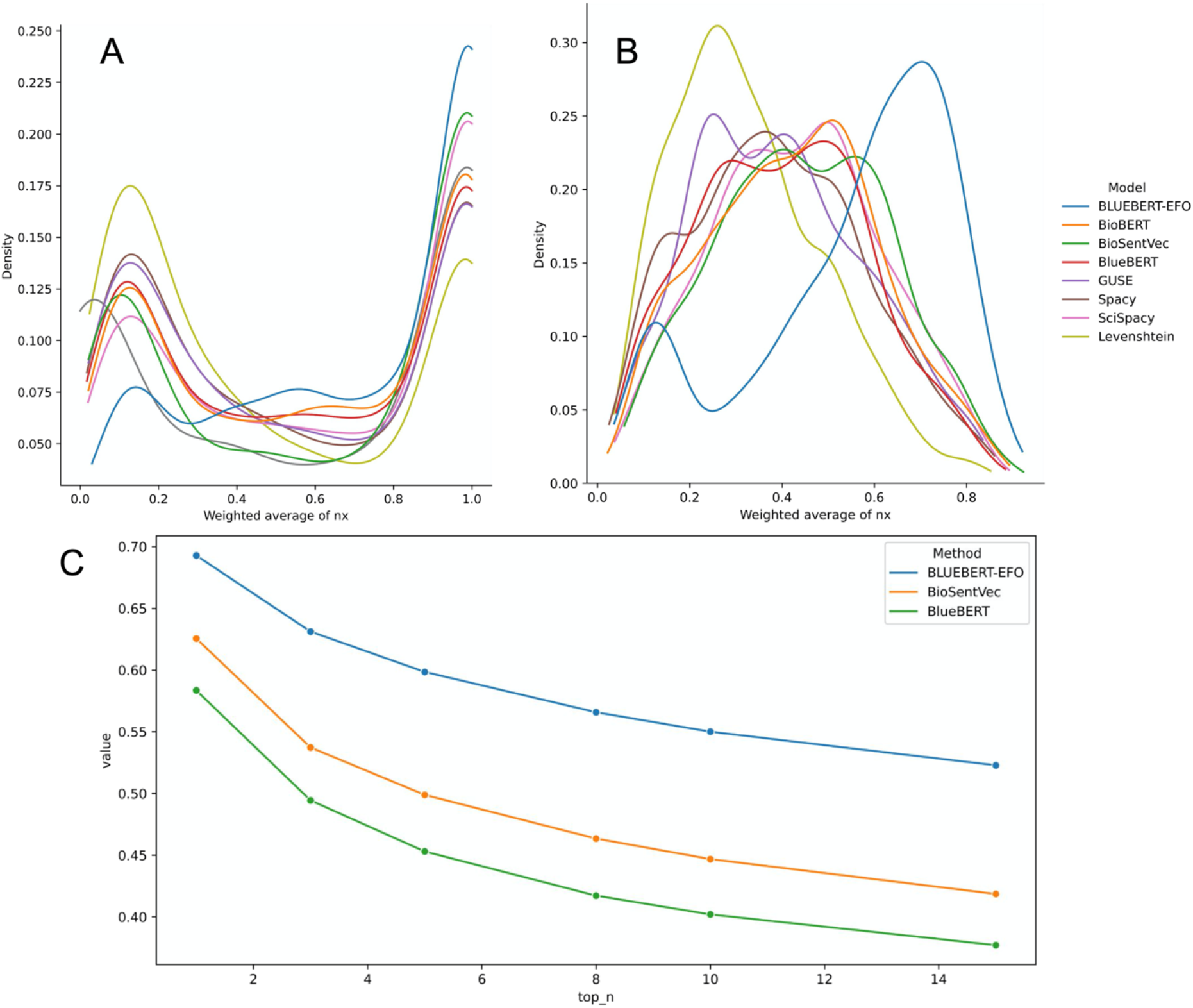
Distribution of predicted EFO-Batet scores by method. (A) Distribution of EFO-Batet score for the highest-ranking (top 1) match for each query term; (B) Distribution of weighted average EFO-Batet score for the top 10 matches for each query term. (C) Averaged sum of the top N weighted averaged EFO-Batet score of the predicted EFO candidates for a query term, for subset methods of BlueBERT-EFO, BlueBERT, and BioSentVec (full results are available in **Supplementary Figure S10**).

We then extended the analysis to consider a set of top results. Figure 3B shows the distribution of the weighted average EFO-Batet scores for the top 10 EFO predictions for each method (see **Supplementary Figure S4** for violin plot and **Supplementary Table S3** for descriptive statistics on the same data). For top 10 predicted EFO terms, we computed the EFO-Batet score vis-a-vis the manual annotation counterpart, then averaged with the ranking weights (i.e. top prediction getting a weighting of 10, second 9 and so on) to show the aggregate ontology relevance of the retrieved candidates. **Figure 3C** shows the averaged sum of the weighted average scores for each top N level to provide an overall measure on the general ontology relevance of the candidate retrieval for a subset of methods (see supplementary figure S10 for full results). The results suggest that BlueBERT-EFO will generally return a set of traits that are more closely associated with the correct part of the EFO ontology compared to other methods, and corroborates with earlier analysis findings that the finetuning of the BlueBERT language model with EFO structure information will notably improve EFO candidate retrieval.

We also investigated on the performance of a hybrid method (BioSentVec-X-BlueBERT-EFO) where BioSentVec is applied in the first stage to select the top X (e.g. 30) candidates, then BlueBERT-EFO is applied in the second stage to select the top N (e.g. 5) candidates, with the aim to improve inference efficiency as transformer models are more computationally expensive than simpler model architectures such as BioSentVec. **Supplementary figures S5-S7** show the weighted average score distribution for top 1, 5, and 10 matches, and supplementary figure S8 show the averaged sum of weighted average scores for generalised top N levels. These results suggest that top matching results produced by the second stage BlueBERT-EFO in the hybrid methods retain the overall behaviour of BlueBERT-EFO, and is robust to the first stage filtering via BioSentVec.

To try and understand why certain traits are challenging to map, and why others are not, we extracted the top UK Biobank queries which were most and least variable in EFO-Batet score between methods. Details of this can be found in **supplementary text S2**.

### Comparison to other approaches for trait-to-trait mapping

Our final set of analyses explores the differences in direct trait-to-trait mapping of the different models. For each model we estimated trait similarity scores between each trait (n=530, see **System and Methods**) and all others (excluding itself). **Figure 4** shows the results of a Spearman rank correlation analysis comparing the matrices of these pairwise trait-mapping scores between each pair of models. The results broadly indicate three clusters of models. One contains the EFO-Batet (manual mapping) and BlueBERT-EFO scores, suggesting again that the BlueBERT-EFO model, as expected, is predicting distances most similar to that which we find in the EFO hierarchy. A second group contains the other BERT models (BioBERT and BlueBERT) highlighting the similarity between those two transformer models. A third group contains the spaCy, ScispaCy and BioSentVec models, which may represent their shared underlying methodology, (i.e. variations of word2vec). Whilst this analysis can’t tell us which method performs “best” at trait-to-trait mapping, it highlights that these models do perform differently at this task, which should be taken into account in the development of future automated trait-to-trait mapping methods.

**Figure 4:**
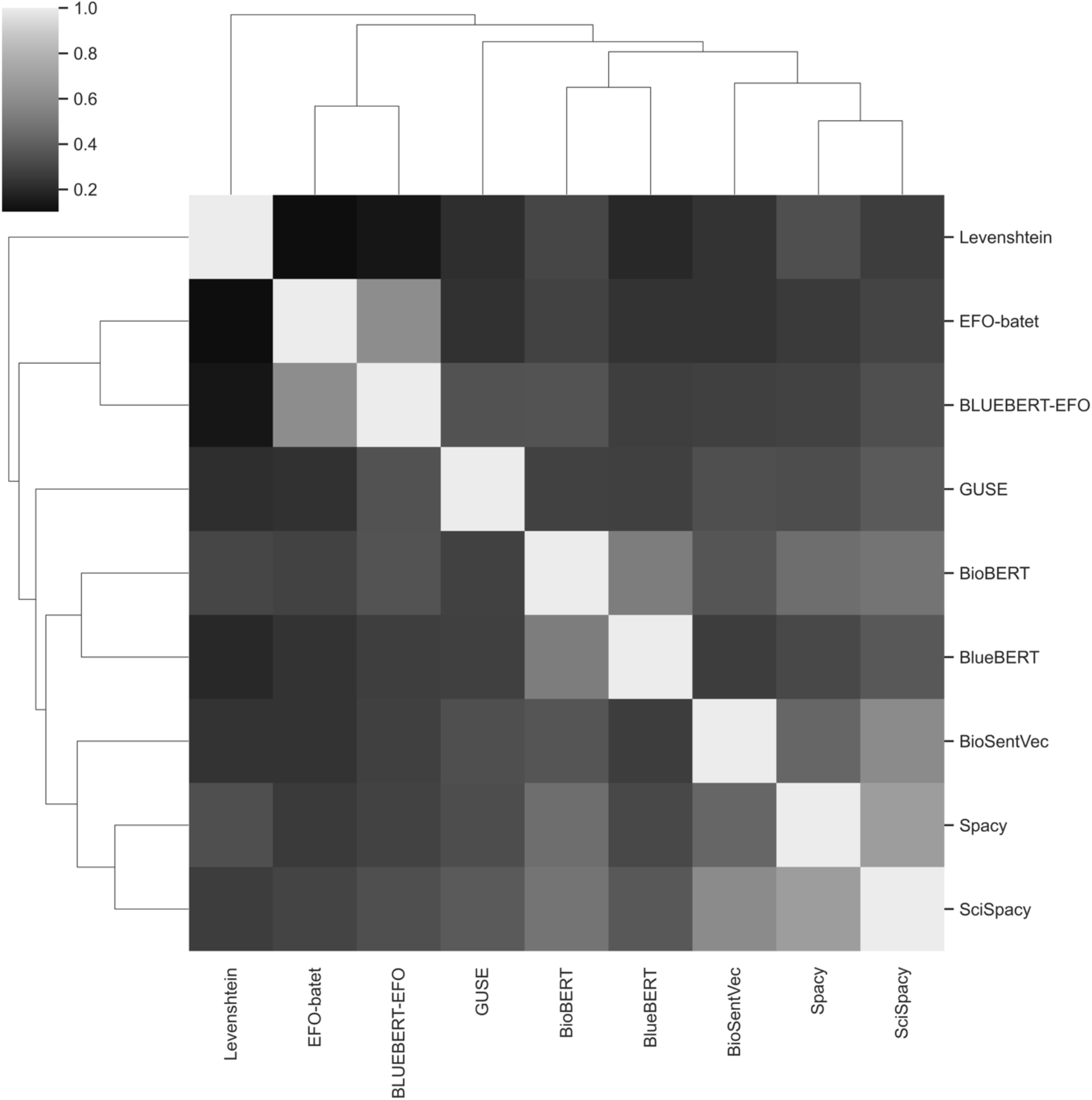
Pairwise plot of spearman correlations between methods based on a matrix of cosine similarity (or equivalent) scores for all pairwise combines of traits (excluding self).

Finally, we present visualizations of the trait similarity scores for all pairwise trait-to-trait mappings for a selected set of 43 traits (representing a mixture of disease, continuous traits and medications) to illustrate how these models perform at this task. **Supplementary Figure S11** is provided as a reference and shows a clustered dendrogram of EFO-Batet scores for the distance between traits in the EFO hierarchy. The clusters represent the relationships between EFO terms as determined by the EFO hierarchy and batet scores. We observe a sharp separation between measurement based quantitative traits and disease traits. This reflects the structure of the EFO, with quantitative traits falling into the “information entity” and disease traits into the “material property” top-level branches of EFO (https://www.ebi.ac.uk/ols/ontologies/efo).

Using the same 43 traits, we then produced a matrix of trait-to-trait distance scores for each model, but this time based on cosine distances (or equivalent – see **System and Methods**). These matrices were compared to each other using the Mantel test^51^, a method to compute correlation distances between matrices (**supplementary Figure 12**). Here we see a similar pattern, with the BlueBERT-EFO and EFO-Batet (i.e. position in the EFO hierarchy) scores clustered together. This similarity is obvious in the BlueBERT-EFO clustermap (**supplementary Figure 13**) where there are some clear differences, but the major distinction between quantitative traits and disease is present, with almost exactly the same traits clustering into the same two groups. This likely reflect the finetuning of this model to EFO.

## Discussion

A number of approaches exist for text matching and semantic representation of text. We set out to investigate the use of these approaches for the automated mapping of human trait names to ontologies (using the specific example of EFO) and explore how they perform at direct trait-to-trait mapping.

### Comparison of approaches for automated mapping to ontology

Our analyses illustrate that using text embeddings to map biomedical variables to EFO has a fairly high error rate, but is at least comparable to existing approaches (e.g. Zooma^14^). Given the ease of use and scalability of some of the models, we recommend this approach when tackling problems that involve many thousands of variables and manual annotation is not feasible. When attempting an exact match (i.e. top match) BioSentVec^21^ appears to perform best in terms of speed, precision and accuracy. However, if it is more important that the top *N* predictions are close to the truth, then BlueBERT-EFO consistently out-performed all other models.

It is important to note that several of the models had similar performance at finding a top match, with the group including BioSentVec, BLUEBERT-EFO, Zooma and ScispaCy^46^ showing little statistical evidence of a difference. It is important to note that the standard Zooma tool also brings the benefit of continually updated manually curated mappings^14^.

Embedding methods appear to perform well when the query string describes a single event or entity, e.g. “whooping cough / pertussis”. They perform poorly when the query string describes multiple entities, e.g. “hiv disease resulting in malignant neoplasms”. This is perhaps not surprising, as the embedding of this phrase is unlikely to be close to either HIV or cancer terms. Addressing such traits therefore remains a complex challenge, i.e. properly identifying mentioned concepts via named entity recognition (NER) and then incorporating pretrained concept embeddings from the knowledge base to the document embeddings ^52,53^. In other words, a complex processing system which includes major components of NER, document level embeddings, and concept embeddings is required to approach mapping of complex traits in a generalised and robust manner, though we are keen to explore this aspect in future research.

We compared our models to a manually mapped set of trait names, but it is important to recognise this may itself contain errors. Supplementary file S7 lists examples where no models predicted an EFO term with an EFO-Batet score >0.95. Here, for example the query term “malignant neoplasm of colon” was manually mapped to “colon carcinoma”. However, six of the models predicted the EFO term “malignant colon neoplasm” which has an EFO-Batet score of 0.86 and is therefore a better fit. (It is possible these differences reflect changes in the EFO since the initial mapping rather than a mapping error).

### Comparison of approaches for trait-to-trait mapping

Mapping traits directly between two datasets has potential value, but in the absence of a benchmark it is hard to validate. We therefore focused on variables that had been mapped to a single EFO term, and then refined that further for closer inspection. The use of clustering methods enabled us to manually inspect groups of traits and describe events that agree with standard biomedical knowledge. Our analyses show that by including topological information from a well established ontology like the EFO, the BlueBERT-EFO model can create sensible pairwise distances between variables, without actually mapping to ontology.

When focusing on a specific set of traits, we see that whilst the finetuning of BlueBERT-EFO has produced a model which reflects major patterns in the EFO hierarchy, there are some differences. One example is the loss of the “angina”, “worrier / anxious feeling” cluster (present in EFO, **Supplementary Figure S11**), with “angina” joining the larger disease cluster next to “atrial fibrilation and flutter” and “worrier / anxious feeling” moving next to “neuroticism score” (**Supplementary Figure S13**). The manual EFO term assigned to “angina” was “EFO_0003913” (angina pectoris, http://www.ebi.ac.uk/efo/EFO_0003913) which can be found within the “material phenotype” EFO group as it is listed as a “Phenotype abnormality” and not a disease. Even though the BLUEBERT-EFO model has been finetuned on the EFO hierarchy, the biomedical literature underpinning the model has created distances placing “angina” with other diseases rather than measurements. This highlights the subtle balance of information contained within this model.

Interestingly, the BlueBERT-EFO model fails to group together the neurological illnesses (“parkinson’s disease”, “alzheimer’s disease” and “secondary parkinsonism”). Looking at the other models, several also fail to do this, often grouping traits with the word “disease” together (**Supplementary Figures S14-20**). However, BioSentVec, BlueBERT and BioBERT appear to group these appropriately. This highlights one of the key challenges that the developers of these models face: how to distinguish between informative words and ignore the generic (e.g. “disease”). This point is again present in the BioBERT cluster map (**Supplementary Figure S19**), with “weight” an outlier to all other traits, suggesting this term was not sufficiently similar to anthropometric traits.

It is worth noting, that the alternative methods to using language models for this type of distance analysis appear to perform less well (e.g. Levenshtein edit distance, **Supplementary Figure S14**). Other established methods, such as Zooma, are just not possible to use when comparing data in this way.

At the moment there is no practical alternative automated approach to trait to trait mapping, so our results using language models are promising. However, they are far from perfect with many cases of traits not grouping together as we might expect, and the models often focusing on generic words such as disease over and above other more defining terms. This approach therefore requires further development before it can be of practical use.

### Use cases of these models

The models are imperfect but are still successful in mapping 40% of trait names in the dataset we used. One obvious use case would be a semi-automated mapping tool which would provide a suggestion for the user to approve or edit. As highlighted above, many simple trait names map well, and it is the more complex traits (e.g. combinations of entities) that would need manual intervention.

Another scenario in which an imperfect one-to-many mapping tool like those presented here may be useful is in a “trait name recommender”. One example of this is our OpenGWAS^54^ recommender, which provides recommended trait matches from amongst thousands of GWAS datasets to enable a user to see other relevant GWAS traits they may be interested in. The OpenGWAS recommender uses a combination of ScispaCy and BlueBERT-EFO to search for the top matching GWAS traits in the semantic embedding vector space and optionally predict the ontology relationships between the query term and the match candidates^55^.

In a follow-up study^56^ we applied ScispaCy and BlueBERT-EFO as an ontology mapper in a hybrid architecture, where a first stage model is used to efficiently filter EFO ontology candidates associated with the query ULMS terms, and in the second stage BlueBERT-EFO is then used to predict the ranking of the top N results (similar to results in supplementary figures S5-S8 where BioSentVec was applied as the first stage model). The retrieval results have shown to be sensible for the systematic analysis on medRxiv submission abstracts, without sacrificing inference performance due to the computationally expensive nature of transformer models whilst retaining relevancy in candidate retrieval.

### Conclusions

We have shown that current text matching and embedding approaches offer some promise in the task of mapping traits to ontologies and to each other. However, the mapping is imperfect and unsuitable for fully automated mapping. Models trained on the biomedical literature perform better than more generalised models. Some trait names present in population health datasets such as UK Biobank are complex and their embeddings are unlikely to be very representative; future work should focus on how to handle such trait names.

## Availability

Code is available at https://github.com/MRCIEU/vectology. This contains general methods, examples and the code used to perform the analyses discussed here.

