## supplementary-materials for "Using language models and ontology topology to perform semantic mapping of traits between biomedical datasets"

### Supplementary: Text

#### S1: Finetuning a transformer model with an ontology alignment task

Here we discuss the design and implementation of the ontology alignment task used to train a transformer pretrained model into a finetuned bespoke ontology classifier model (BlueBERT-EFO). This method of model training for entity mapping is not exclusive to the biomedical domain but can be applied to other domain-specific ontology systems. We aim to train a semantic model that learns the topological structure of an ontology system, i.e. the pairwise distance of ontology terms, and thus is able to produce a score that matches the ontology distance for a pair of biomedical terms (which can be terms in an ontology system but also text labels that are semantically similar to ontology terms). We conduct the training process as a finetuning task using a transformer model that has been pretrained in language modelling tasks in the biomedical domain, and therefore is able to complete the training process within a few epochs without having to train the model parameters from scratch.

An ontology system (e.g. EFO in our case) can be simplified as a graph $G(V, E)$ where a node (vertex) $v_\alpha \in V$ is an ontology concept term (e.g. "body mass index") and a link (edge) $e(v_\alpha,\ v_\beta) \in E$ is the parent-child relationship between $v_\alpha$ and its parent term $v_\beta$ (e.g. "body weights and measures"). The pairwise shortest distance $e'(v_\alpha,\ v_\beta)$ containing multiple $e$ steps between two ontology term nodes can serve as a proxy measure to represent the distance between the two concepts in an ontology, and the structure of the ontology graph can be formulated as the set of $\{\{v_\alpha, v_\beta, e'(v_\alpha, v_\beta)\},\ \{v_\alpha, v_\gamma, e'(v_\alpha, v_\gamma)\},\ \ldots\}$. Therefore a machine learning predictor based on a transformer language model can learn the structure of the ontology graph as a finetuning task and be used to infer the distance of two concept nodes as a bespoke predictor model. In addition, as the transformer language model has been pretrained with the knowledge of semantic entity representations, the inference can be generalised to apply to entity text labels not necessarily identical to the concept terms in $V$ but also terms that are semantically similar to those concepts (e.g. the label term of a disease or a phenotypic trait). We refer to this training task as an ontology alignment task where the language model learns to align the semantic distance of two concept terms as the distance of two ontology concepts in an ontology system. Regarding the training dataset, we construct the pairwise distance of the EFO terms and expand it to include the EFO annotations of phenotypic traits in the EBI GWAS Catalog ("All studies v1.0.2 - with added ontology annotations, GWAS Catalog study accession numbers and genotyping technology" available from[1]) where we set the distance between a GWAS catalog trait and its annotated EFO term to be 1 (we identified 48 overlapping trait terms between this trait extension dataset and UK Biobank mapping dataset in the main analysis where the majority of the mappings are between identical terms e.g trait term "pain" to EFO term "pain", and would be expected to get mapped by the pre-finetuning basis model, and therefore we think the introduction of bias due to overlap is minimal). We then randomly split this dataset into a training sample and a validation sample and evaluate the performance metrics on the validation sample. The training task is implemented as a sequence regression task where the model produces a floating point score given the vector embedding of a tokenised sequence pair (e.g. the pair of "body mass index" and "coronary heart disease") and learns to minimise mean square error of predicted score and the true ontology distance. As the basis transformer model (a “BlueBERT-base uncased (bluebert_pubmed_mimic_uncased_L-12_H-768_A-12)” model where its parameter weights have been pretrained on PubMed abstracts and MIMIC-III clinical notes) has been trained as a language model that achieves satisfactory results[2] in the BLUE benchmark for a range of tasks on language understanding and text mining in the biomedical domain, we are able to produce a BlueBERT-EFO model with good training results (0.99 in explained variance for the validation set) with 3 epochs of finetuning.

#### S2: Traits which are challenging to map

The EBI EFO dataset is very diverse and contains some trait names which we would expect language models to find challenging. To explore how this might impact on the different methods we extracted the top 5 UK Biobank queries which were most variable in EFO-Batet score between methods. We filtered our results to 'Exact' mapping types (**supplementary table S1**), then sorted by the descending standard deviation of EFO-Batet score to identify entries with the most varied EFO predictions (**supplementary file S6**). We also extracted the ranking of the correct manual assignment from the results for each method.

The results of this analysis are shown in **supplementary table S5**. Through each of these queries and their corresponding EFO mappings, we observed some of the idiosyncrasies associated with the different models. For example, Query 1 ("unspecified jaundice") was manually mapped to the EFO term "jaundice". Five of the models also mapped to this, however, the leading word "unspecified" appears to have led the other four models (Levenshtein, GUSE, BioBERT and BlueBERT-EFO) to map to the distant EFO term (EFO-Batet score of 0.09) "obstructive jaundice". In most cases the term "jaundice" was high up in the ranked list of predictions, however this was not the case for BlueBERT-EFO. Query 2 (anaemia) was manually mapped to the EFO term "anemia". Whilst, we might assume the slight spelling change may affect the models, the greater problem seems to be that some models (e.g. ScispaCy, spaCy and GUSE) map to specific subtypes of anemia. This may occur because these models simply have no embedding for the prefix terms, and so are just matching to "anemia" (and arbitrarily selecting from amongst terms which contain "anemia"). This is supported by "anemia" being the second ranked prediction for each. Interestingly, both Zooma and BLUEBERT-EFO ranked "anemia (disease)" at the top suggesting that perhaps the spelling did affect these models. Query 4, "postpolio syndrome" highlights how varied results can be. Many of the models predicted the same EFO term as that manually assigned, but some seemed to focus on the word "syndrome" (perhaps having no encodings for "postpolio") and BlueBERT-EFO predicts a general term "syndromic disease".

In addition to traits with high heterogeneity between models, we also analysed cases where all traits mapped with high accuracy, and where all traits mapped with low accuracy (**supplementary files S7 and S8**). In **supplementary file S8** we can see 79 cases where no model predicted an EFO term with an EFO-Batet score > 0.95. Many of these are complex trait names, but interestingly some such as "high cholesterol" represent cases where we would expect a model to predict the same EFO term as that manually assigned (hypercholesterolemia), but instead most predicted "cholesterol" first.

### Supplementary: Figures

**Figure S1:** Number of matching predictions by MAPPING_TYPE with an EFO-Batet score >0.9. The Total bar contains all manual mappings, subdivided into Exact, Broad (parent term), Narrow (child term) and Other. Each other bar represents the number of traits exactly matched by the named method to the manual mapping for that trait, with the same subdivisions.

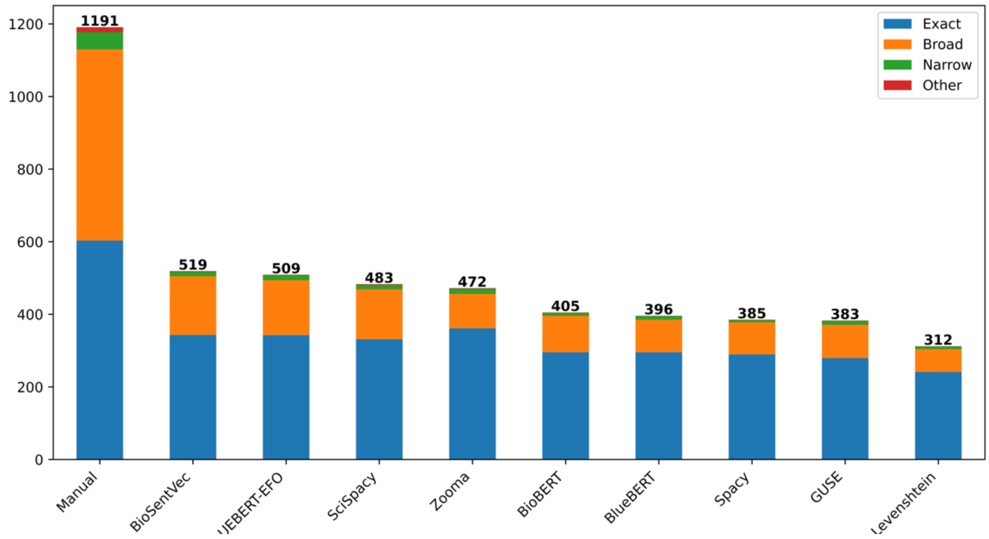

**Figure S2:** Number of matching predictions by MAPPING_TYPE with an EFO-Batet score >0.8. The Total bar contains all manual mappings, subdivided into Exact, Broad (parent term), Narrow (child term) and Other. Each other bar represents the number of traits exactly matched by the named method to the manual mapping for that trait, with the same subdivisions.

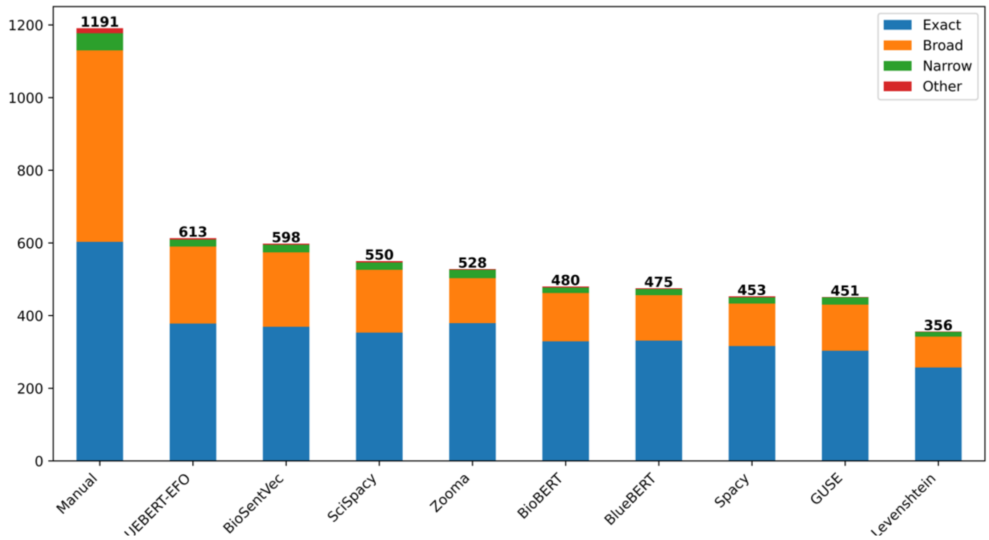

**Figure S3:** Number of matching predictions by MAPPING_TYPE with an EFO-Batet score >0.7. The Total bar contains all manual mappings, subdivided into Exact, Broad (parent term), Narrow (child term) and Other. Each other bar represents the number of traits exactly matched by the named method to the manual mapping for that trait, with the same subdivisions.

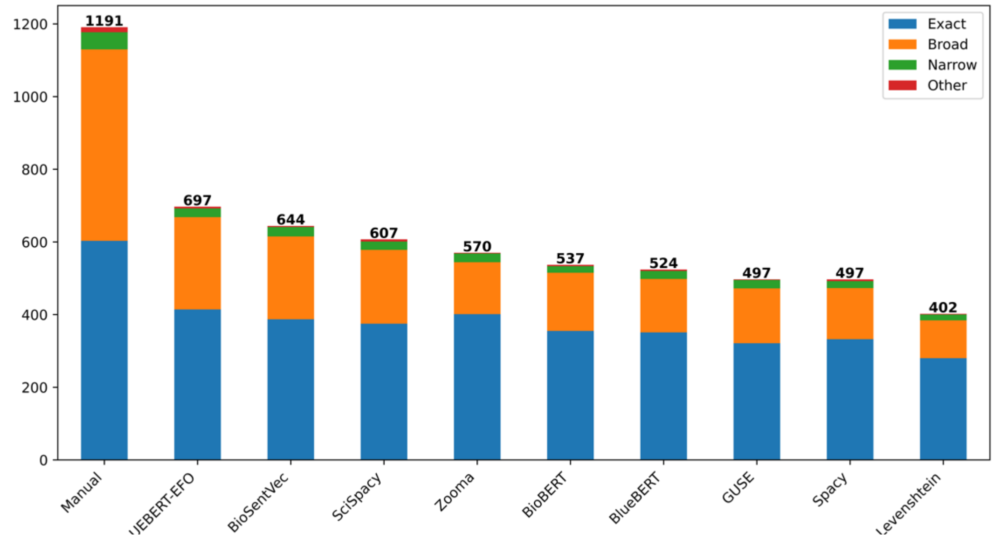

**Figure S4:** Violin plot of weighted average of top 10 EFO-Batet scores for all predictions and models.

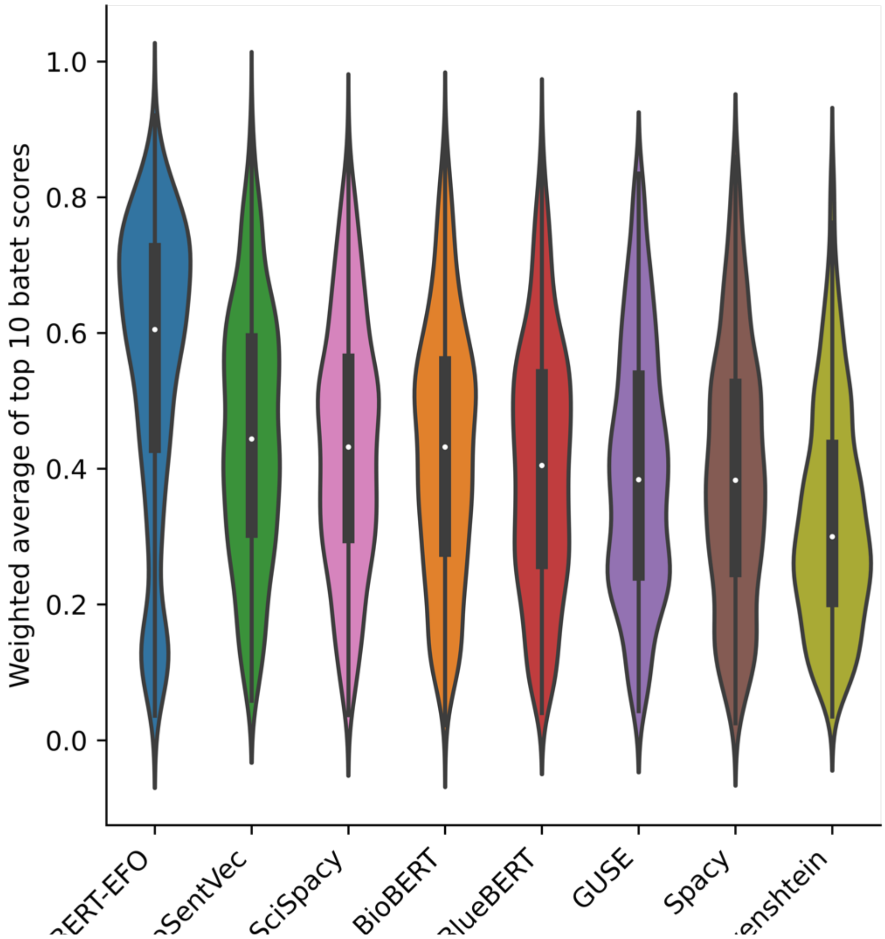

**Figure S5:** Density plots of EFO-Batet scores for hybrid methods, top-1 matches. The hybrid method notation "BioSentVec-X-BLUEBERT-EFO" means in the first stage BioSentVec is used to perform top-X matches in order to filter the list of candidates into a smaller subset, then in the second stage BlueBERT-EFO is then used to perform top-1 matches from the subset to improve inference time.

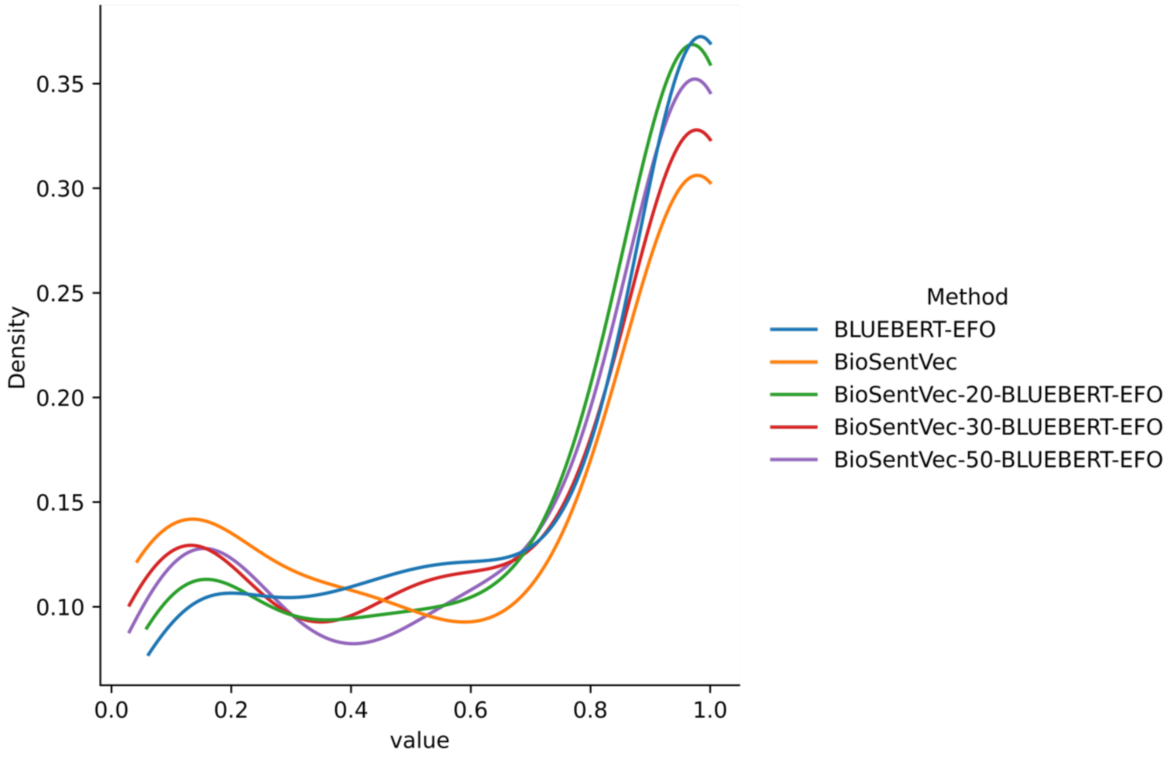

**Figure S6:** Density plots of EFO-Batet scores for hybrid methods, top-5 matches. The hybrid method notation “BioSentVec-X-BLUEBERT-EFO” means in the first stage BioSentVec is used to perform top-X matches in order to filter the list of candidates into a smaller subset, then in the second stage BlueBERT-EFO is then used to perform top-5 matches from the subset to improve inference time.

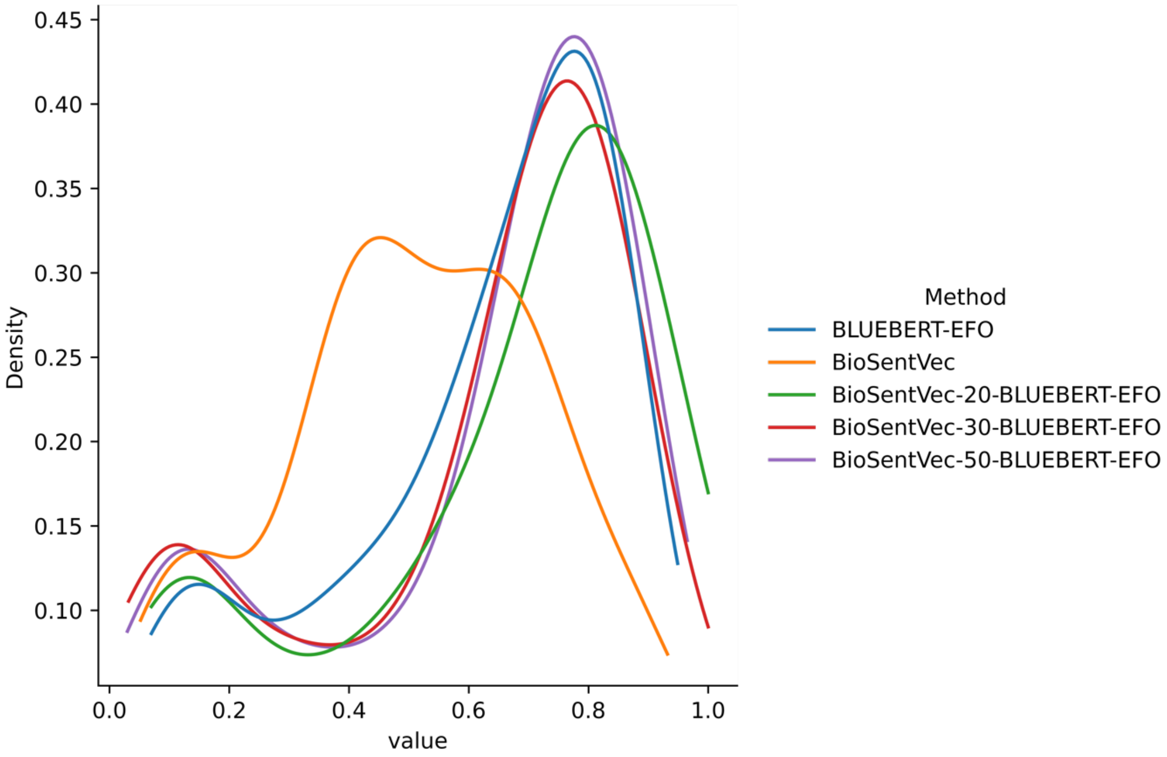

**Figure S7:** Density plots of EFO-Batet scores for hybrid methods, top-10 matches. The hybrid method notation "BioSentVec-X-BLUEBERT-EFO" means in the first stage BioSentVec is used to perform top-X matches in order to filter the list of candidates into a smaller subset, then in the second stage BlueBERT-EFO is then used to perform top-10 matches from the subset to improve inference time.

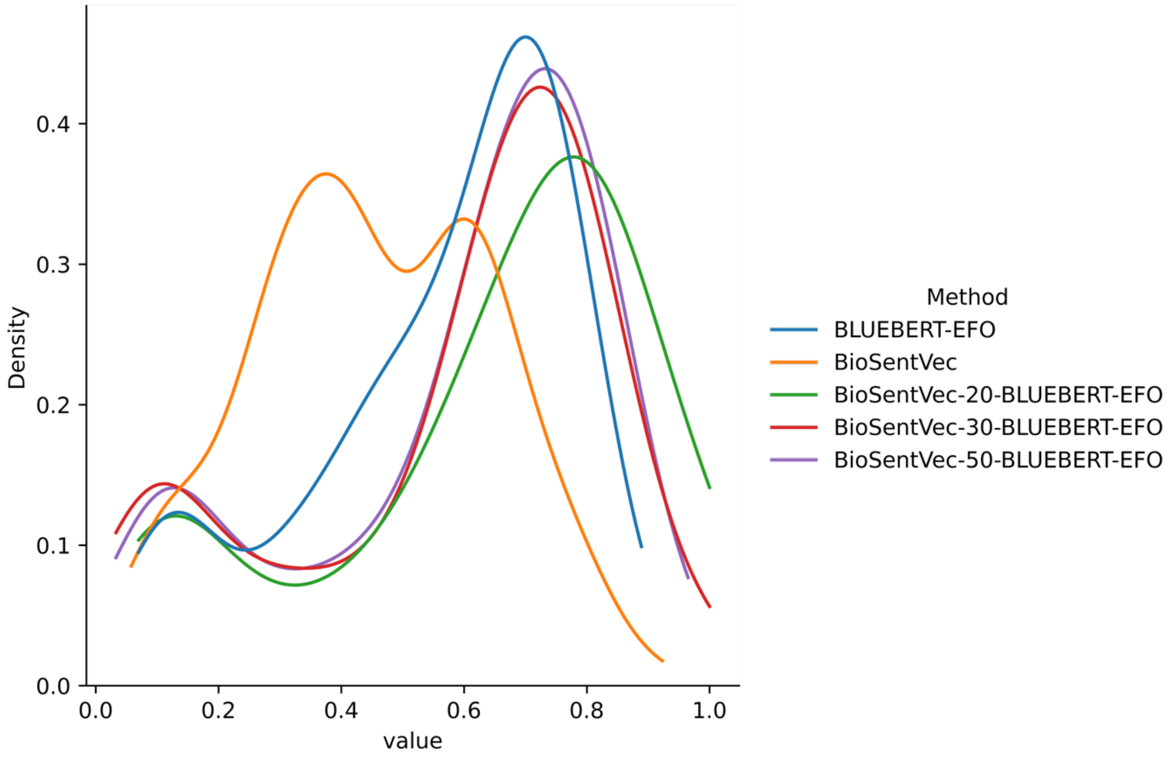

**Figure S8:** Averaged sum of the weighted averaged EFO-Batet score for hybrid methods, by top-N matches. Each point refers to the averaged sum of top-N weighted average EFO-Batet score from a method.

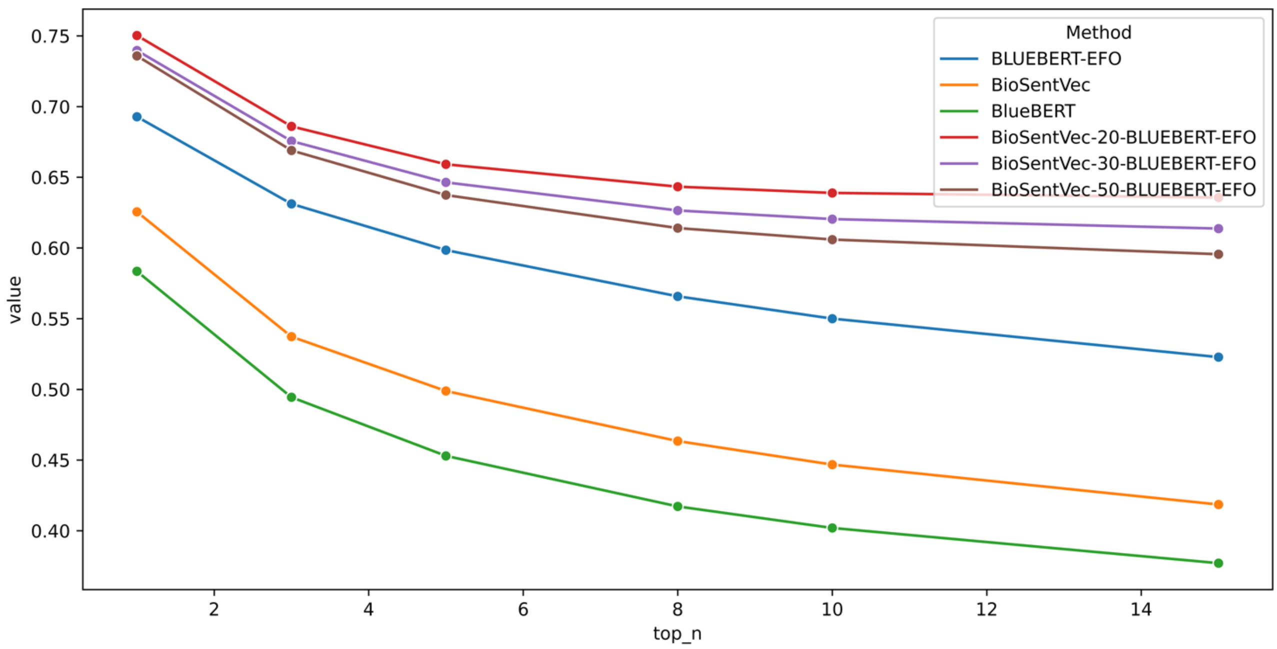

**Figure S9:** Total number of top matching predictions that are equal or above an EFO-Batet threshold. This figure shows the full results in Figure 2B.

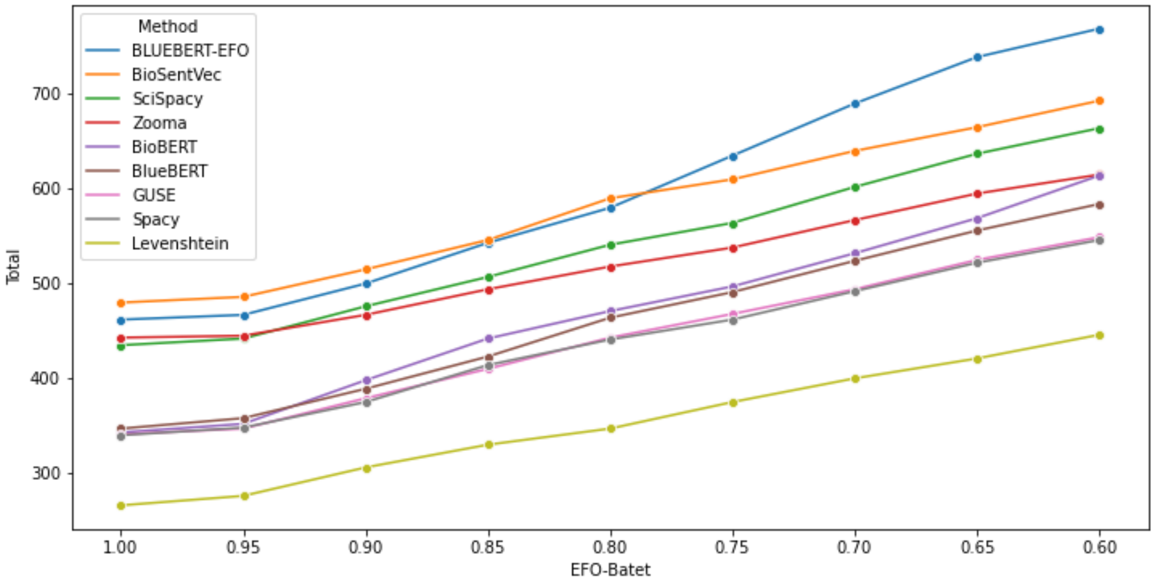

**Figure S10:** Averaged sum of weighted averaged EFO-Batet score of the predicted EFO candidates for a query term. This figure shows the full results in Figure 3C.

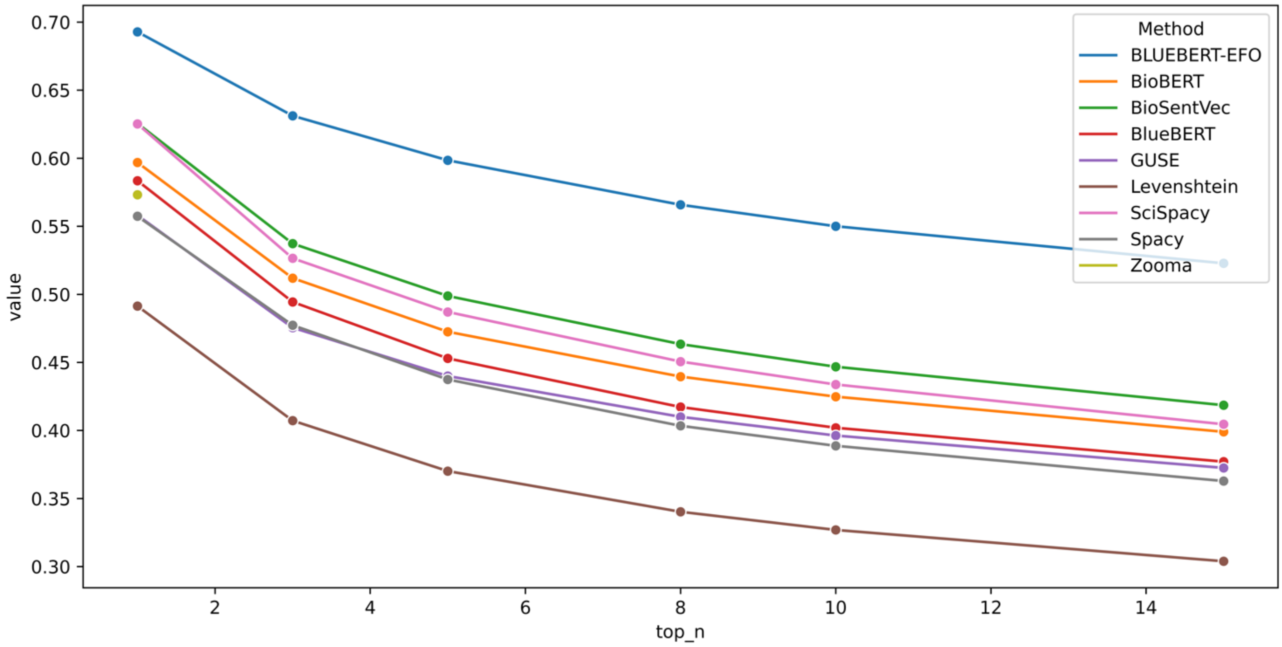

**Figure S11:** Clustered dendrogram of pairwise EFO-Batet scores based on position in the EFO hierarchy.

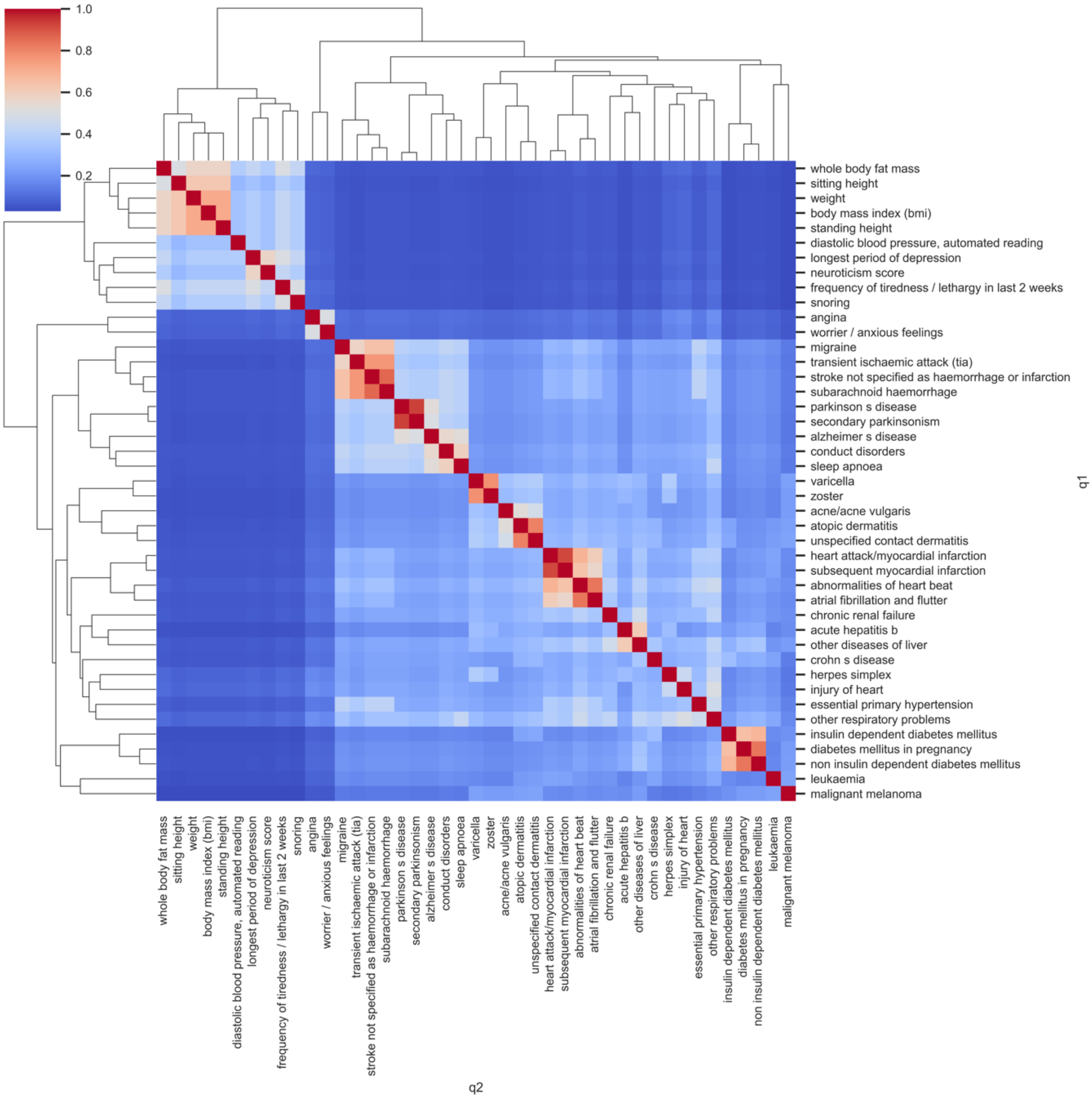

**Figure S12:** Pairwise plot of mantel scores for manual sample.

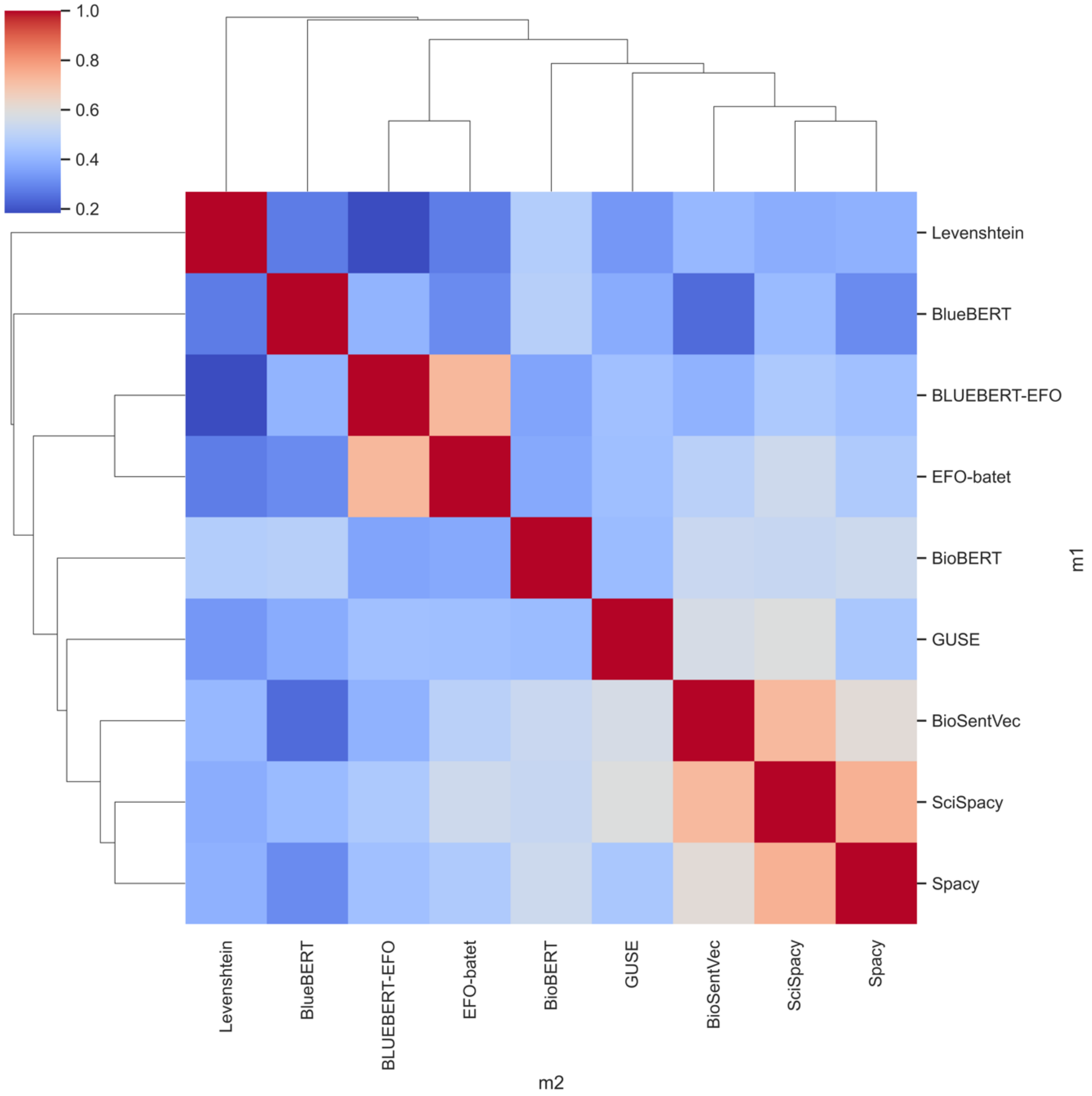

**Figure S13:** Clustered dendrogram of pairwise BLUEBERT-EFO trait similarity scores.

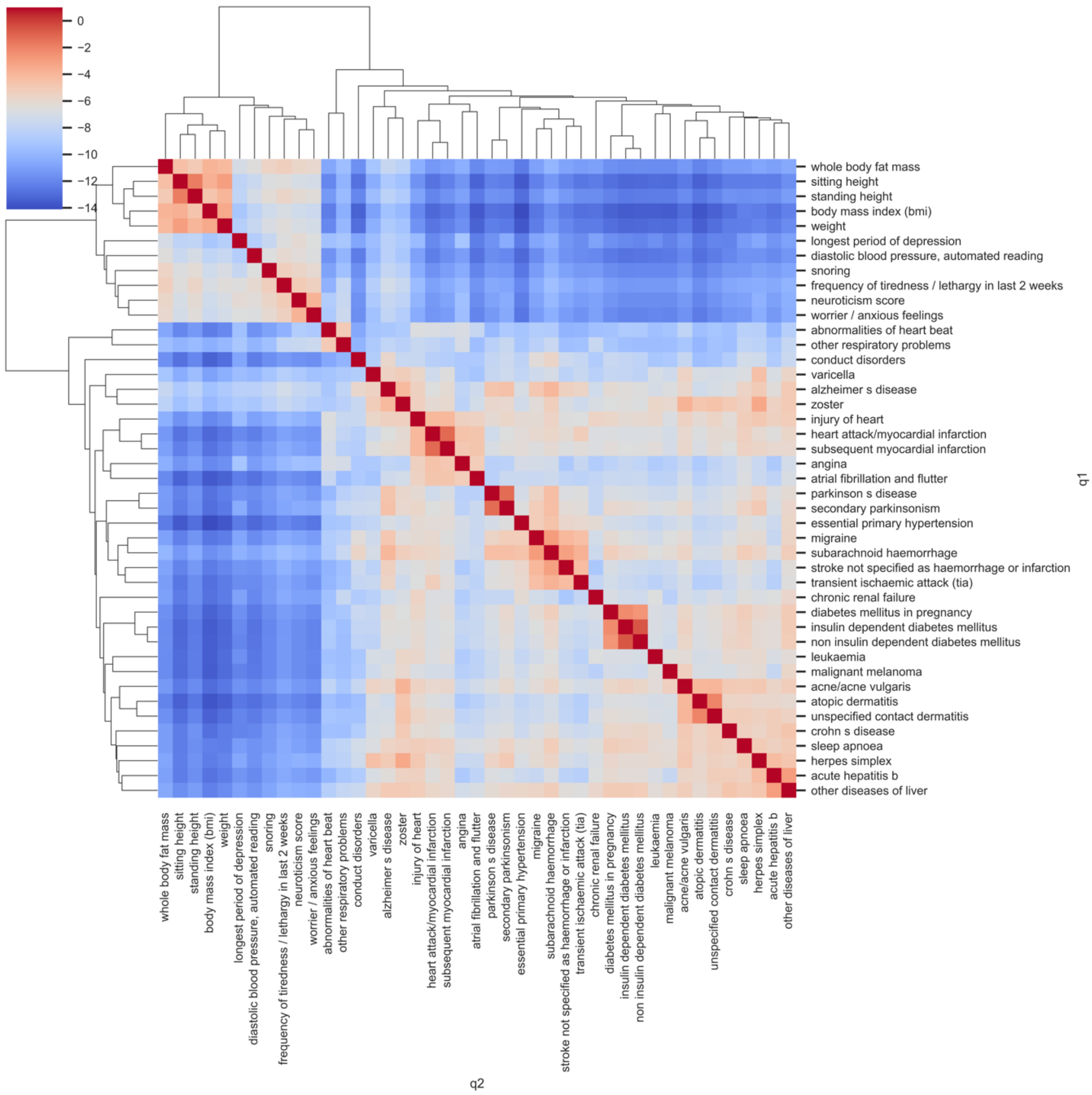

**Figure S14:** Clustered dendrogram of Levenshtein trait similarity scores for manual sample.

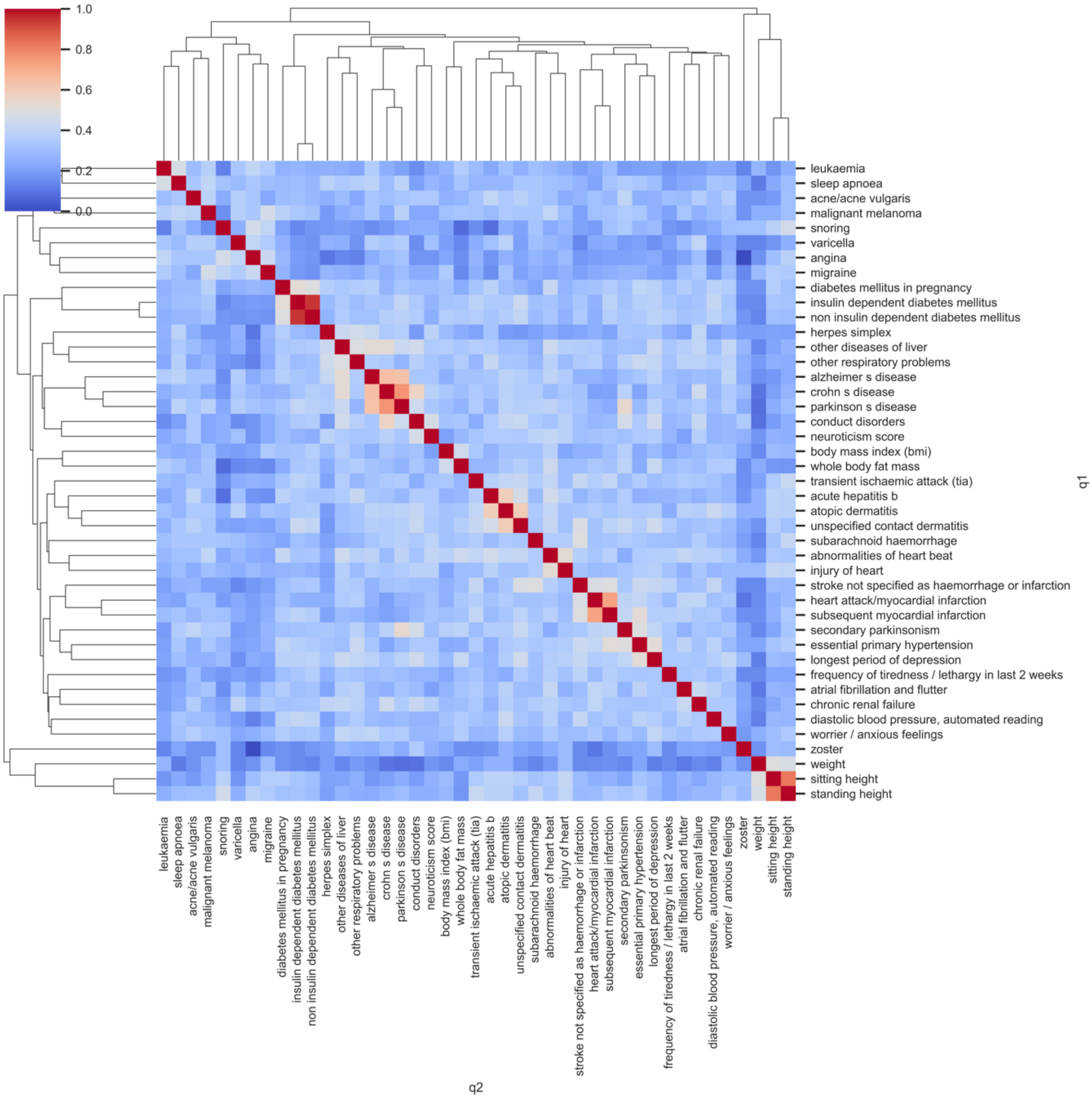

**Figure S15:** Clustered dendrogram of ScispaCy trait similarity scores for manual sample.

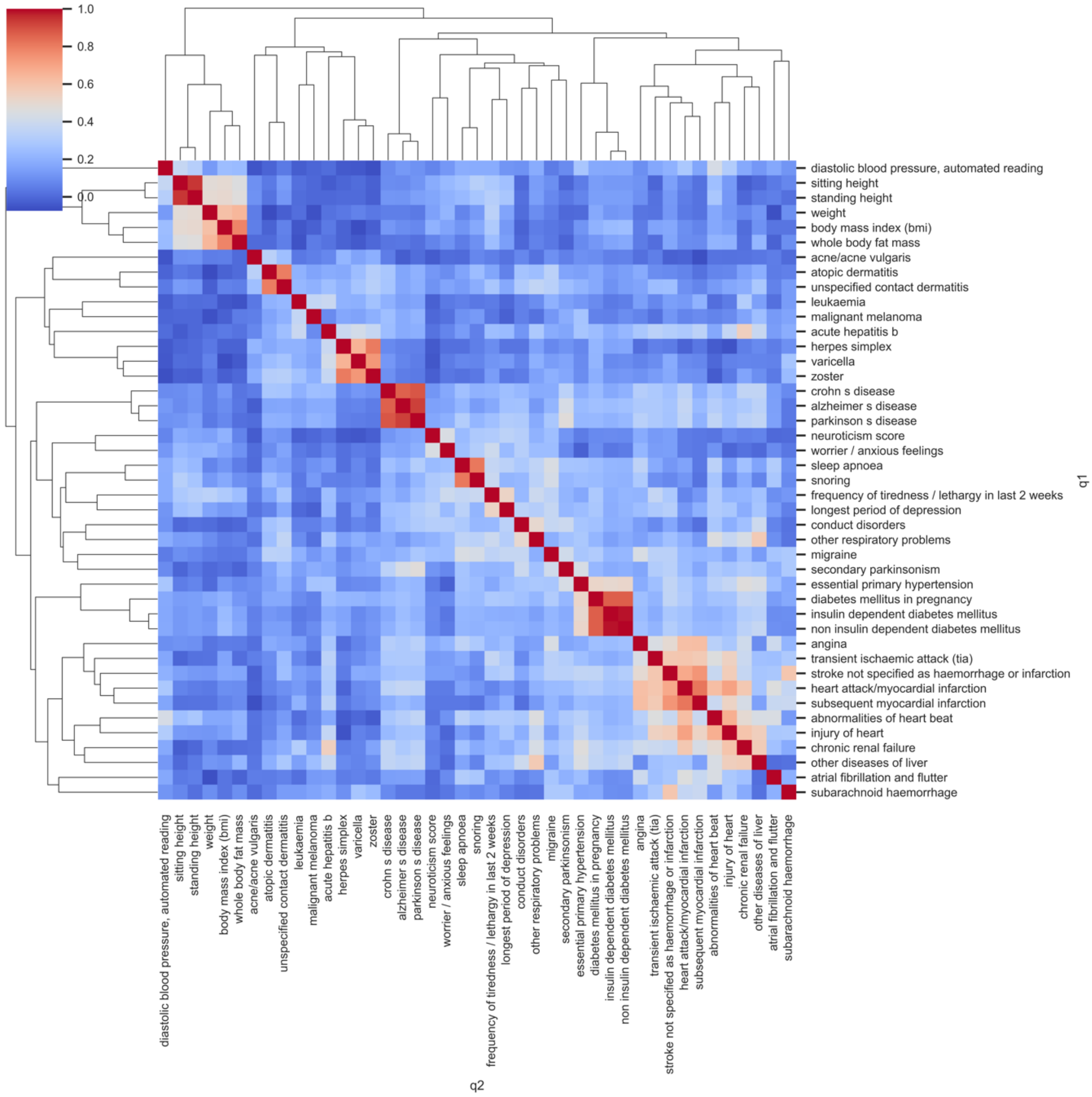

**Figure S16:** Clustered dendrogram of spaCy trait similarity scores for manual sample.

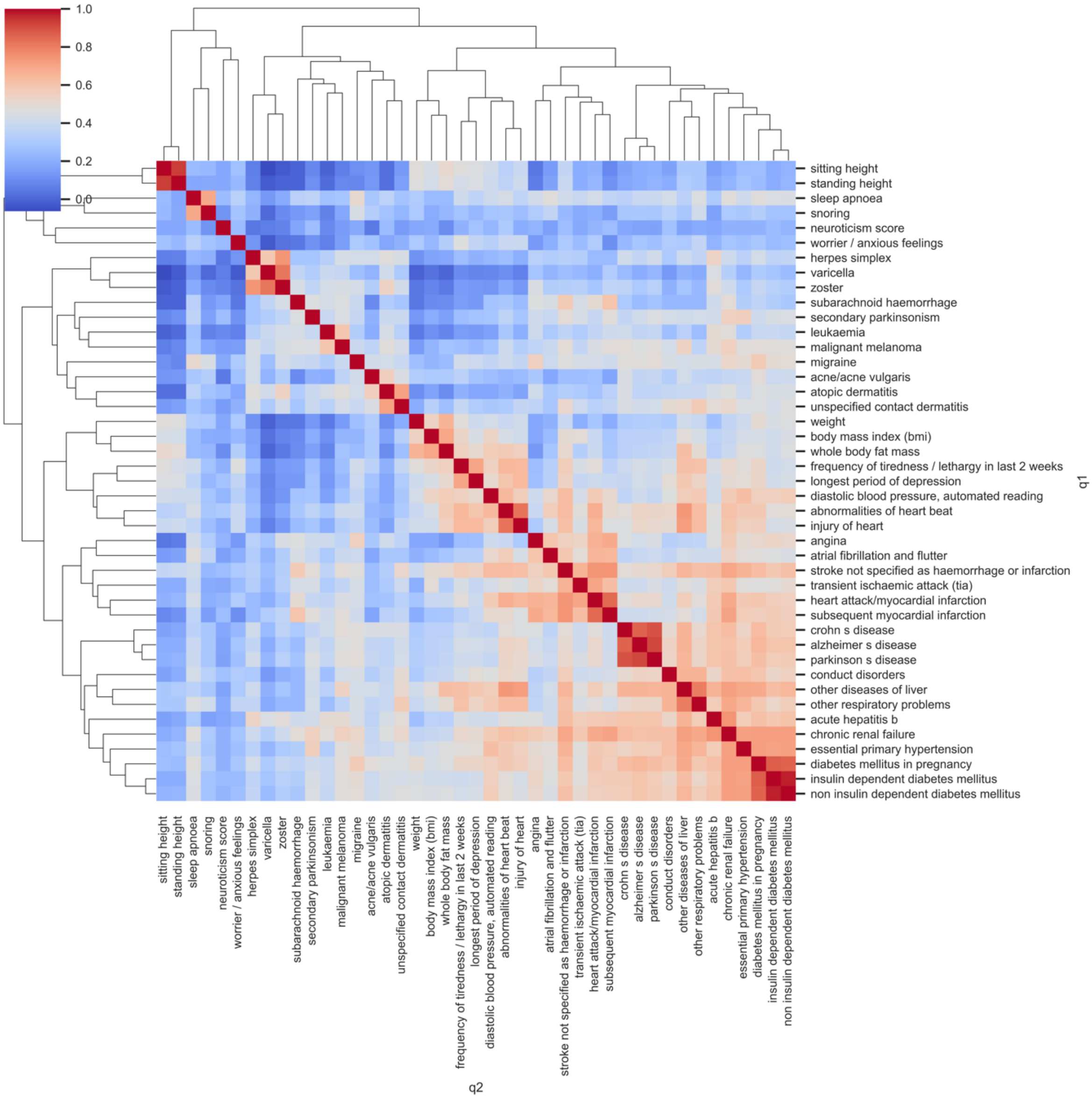

**Figure S17:** Clustered dendrogram of GUSE trait similarity scores for manual sample.

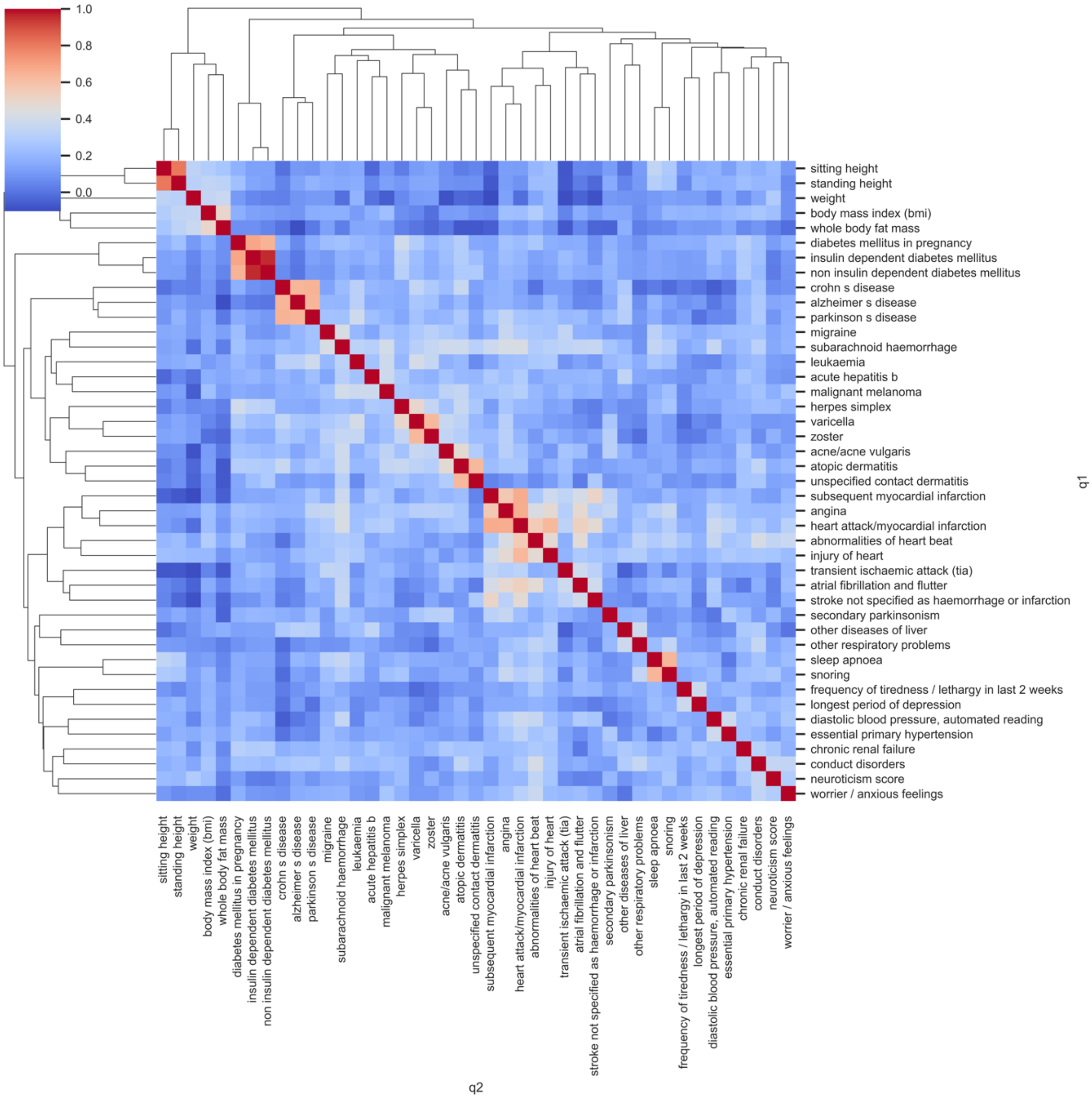

**Figure S18:** Clustered dendrogram of BlueBERT trait similarity scores for manual sample.

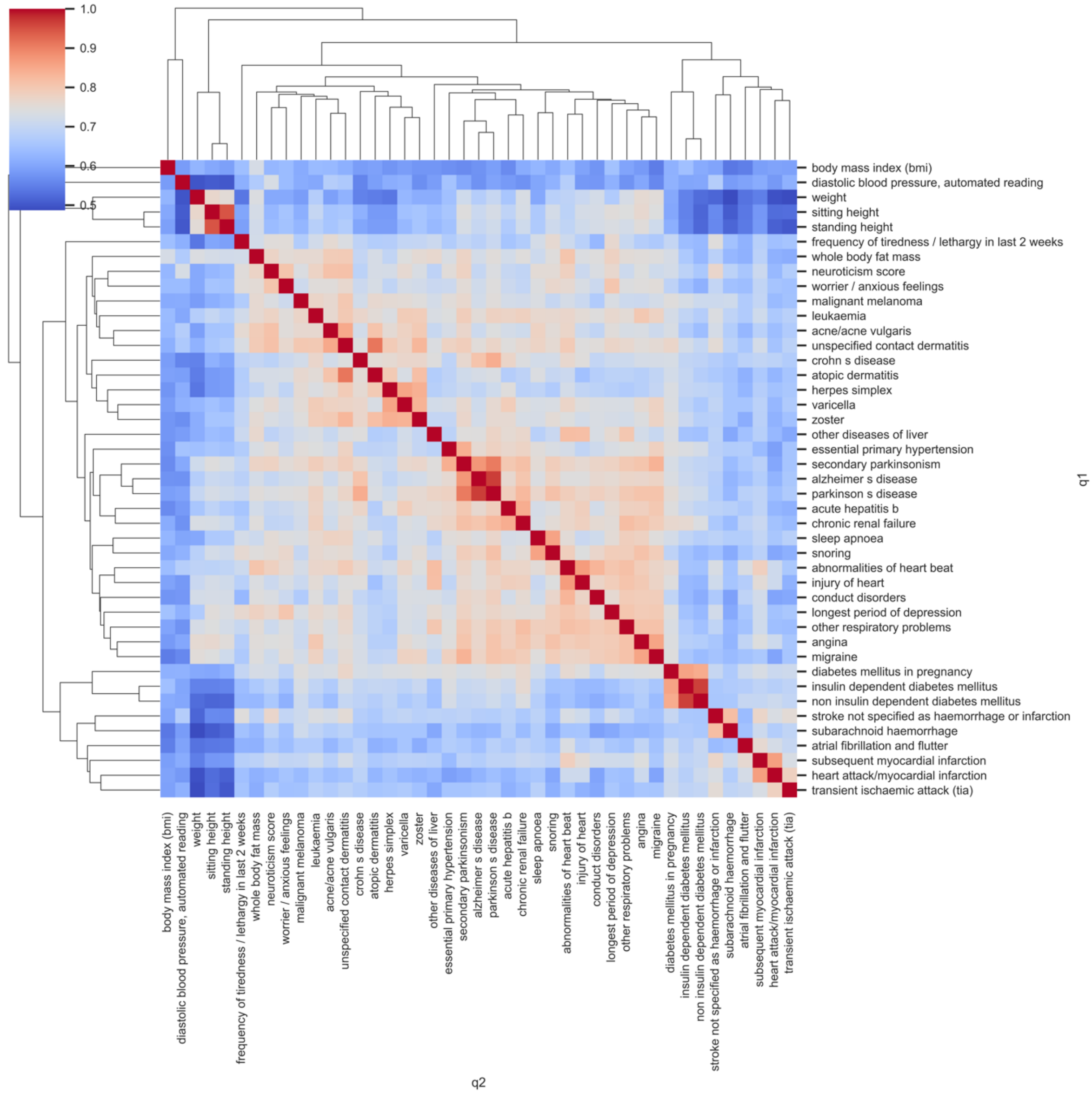

**Figure S19:** Clustered dendrogram of BioBERT trait similarity scores for manual sample.

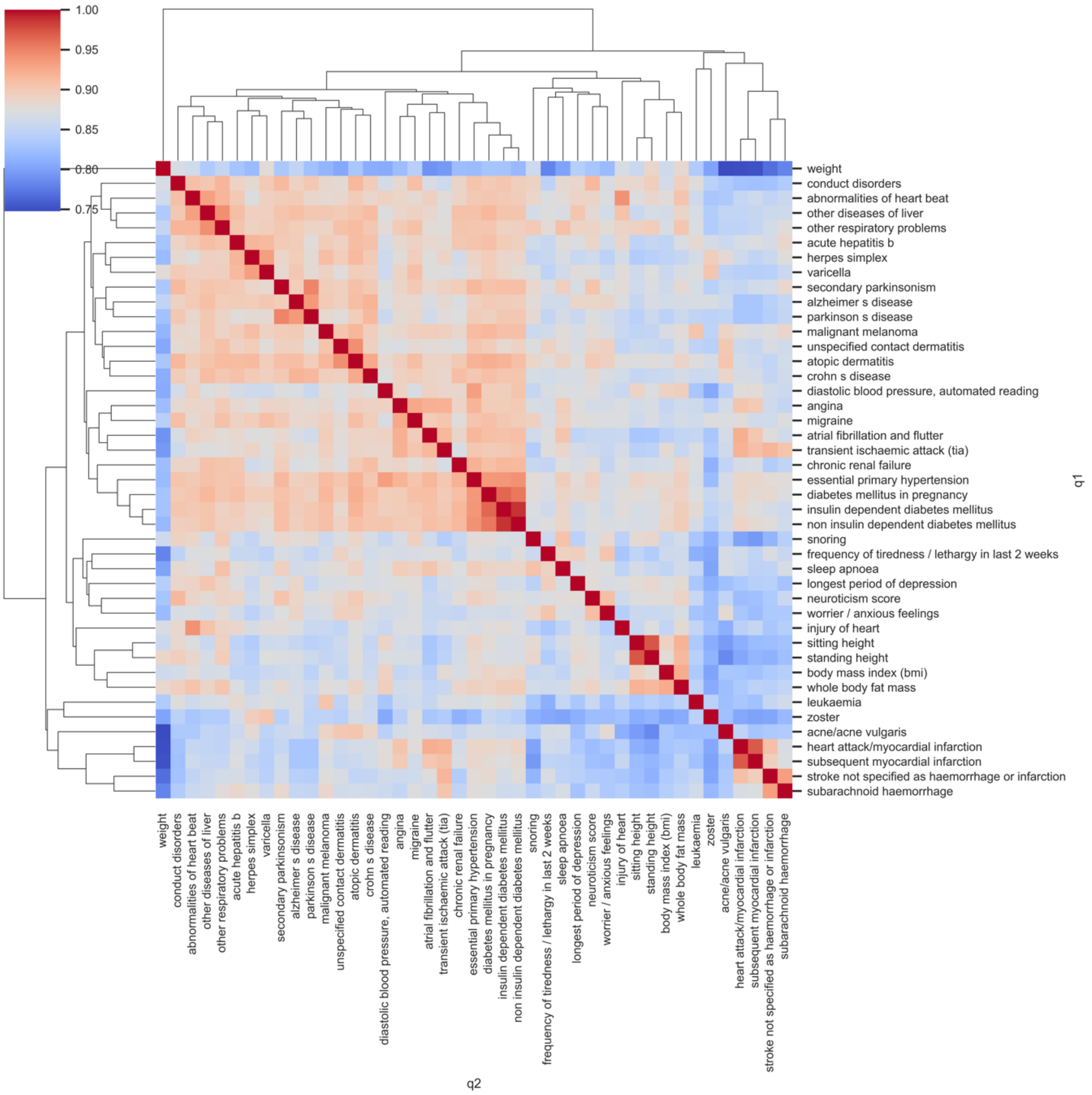

**Figure S20:** Clustered dendrogram of BioSentVec trait similarity scores for manual sample.

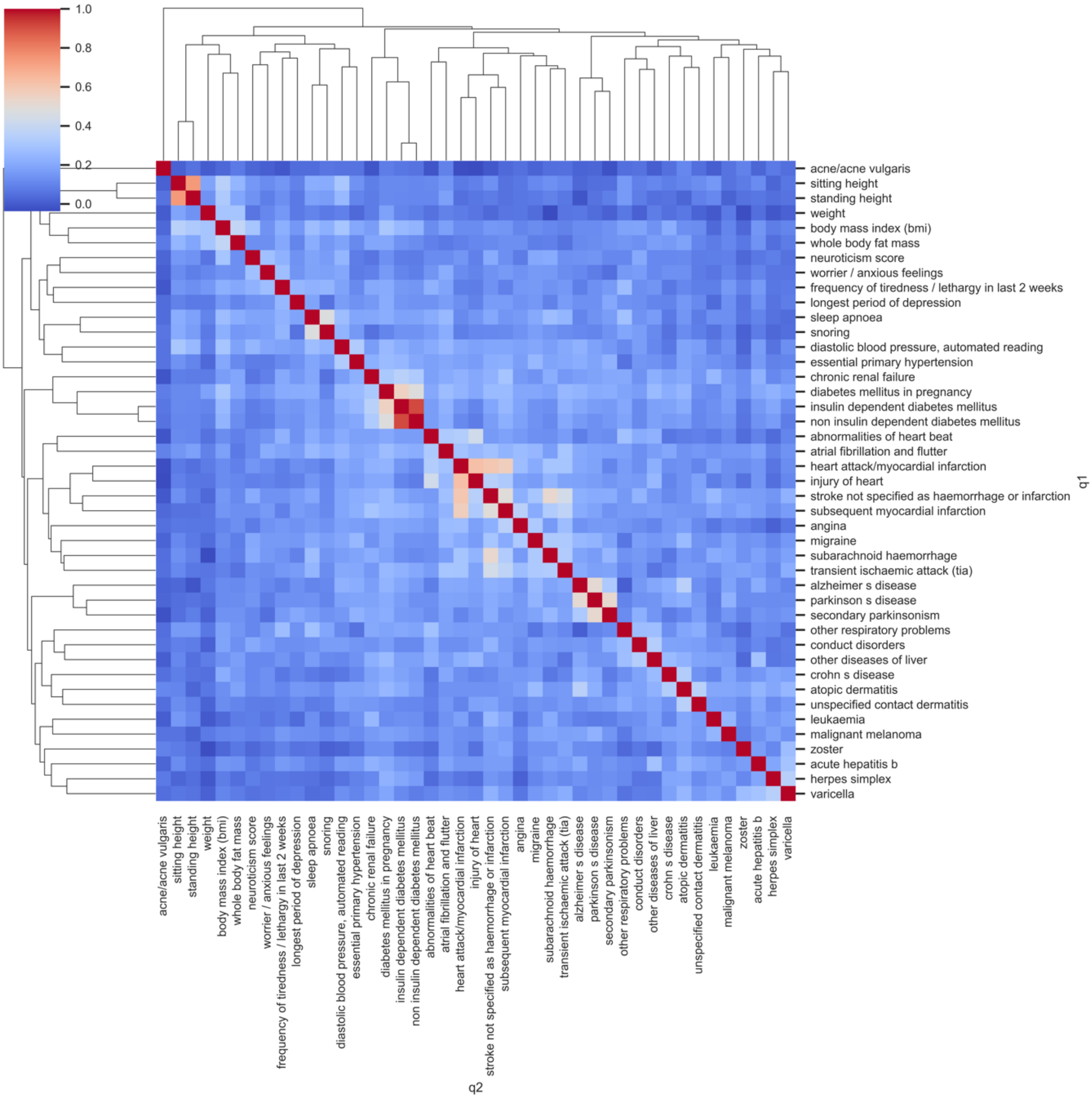

### Supplementary: Tables

**Table S1:** EBI-EFO mapping type names, descriptions and counts.

| Mapping Type | Description | Count |
|---|---|---|
| Exact | Trait and mapped-to term are deemed fully equivalent to one another in definition | 607 |
| Broad | Mapped-to term is a larger concept than the trait itself. The mapped-to trait may be an umbrella concept covering the query trait | 530 |
| Narrow | Mapped-to term is a smaller concept than the trait itself. The query trait may be an umbrella concept covering the mapped-to term | 48 |
| Other | Suggested mapping is either considered too broad, too narrow or incorrect | 14 |

**Table S2:** Summary of methods and models used to map EFO terms to variable text.

| Model/Method | Training data | Dimensions | Distance method |
|---|---|---|---|
| Levenshtein[3] | NA | NA | String matching (edit distance) |
| Zooma[4] | Ontologies and previous mappings | NA | String matching and lookup |
| BioSentVec[5] | PubMed and MIMIC III Clinical notes | 700 | Cosine |
| BLUEBERT[2] | PubMed and MIMIC III Clinical notes | 768 | Cosine |
| BioBERT[6] | Wikipedia, PubMed and PubMed Central | 768 | Cosine |
| GUSE[7] | Wikipedia, etc. | 512 | Cosine |
| spaCy[8] | OntoNotes 5 | 300 | Cosine |

| Model/Method | Training data | Dimensions | Distance method |
|---|---|---|---|
| ScispaCy[9] | PubMed, MesH, Genia, etc | 200 | Cosine |
| BLUEBERT-EFO | EFO | NA | Inferred semantic distance |

**Table S3:** Descriptive statistics of weighted average of top 10 EFO-Batet scores for all predictions and models .

| Model | Mean | Std | Min | 25% | 50% | 75% | Max |
|---|---|---|---|---|---|---|---|
| BLUEBERT-EFO | **0.550409** | **0.218642** | **0.036** | **0.43325** | **0.6055** | **0.72475** | 0.921 |
| BioBERT | 0.425172 | 0.187452 | 0.022 | 0.27925 | 0.4325 | 0.55700 | 0.893 |
| BioSentVec | 0.447244 | 0.187542 | 0.058 | 0.30700 | 0.4450 | 0.59100 | **0.923** |
| BlueBERT | 0.402374 | 0.185323 | 0.040 | 0.26125 | 0.4050 | 0.53875 | 0.884 |
| GUSE | 0.396570 | 0.185861 | 0.043 | 0.24400 | 0.3840 | 0.53600 | 0.835 |
| Levenshtein | 0.326893 | 0.164039 | 0.035 | 0.20500 | 0.3005 | 0.43350 | 0.852 |
| ScispaCy | 0.433470 | 0.183741 | 0.037 | 0.29825 | 0.4320 | 0.56050 | 0.892 |
| spacy | 0.388220 | 0.188961 | 0.025 | 0.24800 | 0.3830 | 0.52225 | 0.860 |

**Table S4:** Test results on statistical difference of two series. Results for Proportions Z-Test report test results regarding statistical difference between the proportions of two top matches (Figure 2). Results for Kolmogorov-Smirnov Test report test results regarding statistical difference between the two series of top EFO-Batet scores (Figure 3). Test statistics are reported outside of parentheses and P-values are reported in parentheses. ***: P-value $\leq 0.01$; **: P-value $\leq 0.05$; *: P-value $\leq 0.1$.

| Model A | Model B | Proportions Z-Test | Kolmogorov-Smirnov Test |
|---|---|---|---|
| BLUEBERT-EFO | BioBERT | 5.155*** (2.540e-07) | 0.143*** (5.384e-11) |
| BLUEBERT-EFO | BioSentVec | -0.754 (4.507e-01) | 0.136*** (5.076e-10) |
| BLUEBERT-EFO | BlueBERT | 4.975*** (6.510e-07) | 0.170*** (1.611e-15) |
| BLUEBERT-EFO | GUSE | 5.245*** (1.567e-07) | 0.208*** (5.229e-23) |
| BLUEBERT-EFO | spacy | 5.290*** (1.226e-07) | 0.204*** (4.230e-22) |
| BLUEBERT-EFO | ScispaCy | 1.142 (2.536e-01) | 0.130*** (3.305e-09) |
| BLUEBERT-EFO | Zooma | 0.844 (3.984e-01) | 0.203*** (6.391e-22) |
| BLUEBERT-EFO | Levenshtein | 8.718*** (2.843e-18) | 0.296*** (1.559e-46) |
| BioBERT | BioSentVec | -5.903*** (3.571e-09) | 0.115*** (2.784e-07) |
| BioBERT | BlueBERT | -0.181 (8.566e-01) | 0.043 (2.249e-01) |
| BioBERT | GUSE | 0.091 (9.278e-01) | 0.075*** (2.577e-03) |
| BioBERT | spaCy | 0.136 (8.919e-01) | 0.076*** (2.217e-03) |
| BioBERT | ScispaCy | -4.019*** (5.833e-05) | 0.077*** (1.633e-03) |
| BioBERT | Zooma | -4.315*** (1.593e-05) | 0.175*** (1.981e-16) |
| BioBERT | Levenshtein | 3.617*** (2.985e-04) | 0.165*** (1.717e-14) |
| BioSentVec | BlueBERT | 5.724*** (1.040e-08) | 0.114*** (3.505e-07) |
| BioSentVec | GUSE | 5.993*** (2.066e-09) | 0.125*** (1.541e-08) |
| BioSentVec | spaCy | 6.037*** (1.566e-09) | 0.127*** (9.286e-09) |
| BioSentVec | ScispaCy | 1.896* (5.801e-02) | 0.055* (5.158e-02) |
| BioSentVec | Zooma | 1.598 (1.099e-01) | 0.170*** (1.611e-15) |
| BioSentVec | Levenshtein | 9.454*** (3.256e-21) | 0.212*** (6.188e-24) |
| BlueBERT | GUSE | 0.271 (7.862e-01) | 0.043 (2.249e-01) |
| BlueBERT | spaCy | 0.317 (7.516e-01) | 0.050* (9.733e-02) |
| BlueBERT | ScispaCy | -3.840*** (1.232e-04) | 0.077*** (1.633e-03) |

| | | | |
|---|---|---|---|
| BlueBERT | Zooma | -4.136*** (3.539e-05) | 0.170*** (1.611e-15) |
| BlueBERT | Levenshtein | 3.796*** (1.469e-04) | 0.133*** (1.496e-09) |
| GUSE | spaCy | 0.045 (9.638e-01) | 0.019 (9.795e-01) |
| GUSE | ScispaCy | -4.110*** (3.964e-05) | 0.099*** (1.650e-05) |
| GUSE | Zooma | -4.405*** (1.056e-05) | 0.170*** (1.611e-15) |
| GUSE | Levenshtein | 3.526*** (4.212e-04) | 0.099*** (1.650e-05) |
| spaCy | ScispaCy | -4.155*** (3.258e-05) | 0.102*** (9.021e-06) |
| spaCy | Zooma | -4.450*** (8.568e-06) | 0.170*** (1.611e-15) |
| spaCy | Levenshtein | 3.481*** (4.989e-04) | 0.099*** (1.650e-05) |
| ScispaCy | Zooma | -0.297 (7.662e-01) | 0.170*** (1.611e-15) |
| ScispaCy | Levenshtein | 7.598*** (3.001e-14) | 0.192*** (1.167e-19) |
| Zooma | Levenshtein | 7.890*** (3.017e-15) | 0.170*** (1.611e-15) |

**Table S5:** The top 5 UK Biobank queries with the largest standard deviation of EFO-Batet scores. These represent the predictions that varied most across the models regarding their location in the EFO compared to the manual prediction. Position of correct prediction is in parentheses (>100 means not in the top 100 predictions) and EFO-Batet score for top prediction is in square brackets. * Query used. ** Correct manual mapping.

| **Model/Method** | **Query 1** | **Query 2** | **Query 3** | **Query 4** | **Query 5** |
|---|---|---|---|---|---|
| UK Biobank trait (Query) * | unspecified jaundice | anaemia | fluid intelligence score | postpolio syndrome | secondary parkinsonism |
| Manual EFO mapping (EBI EFO dataset) ** | jaundice | anemia | intelligence | postpoliomyelitis syndrome | secondary Parkinson disease |
| Levenshtein | obstructive jaundice (5) [0.09] | anemia (1) [1.0] | intelligence (1) [1.0] | postpoliomyelitis syndrome (1) [1.0] | secondary parkinson disease (1) [1.0] |
| Zooma | jaundice (1) [1.0] | anemia (disease) (>100) [0.15] | intelligence (1) [1.0] | postpoliomyelitis syndrome (1) [1.0] | secondary parkinson disease (1) [1.0] |
| ScispaCy | jaundice (1) [1.0] | blackfan-diamond anemia (2) [0.04] | intelligence (1) [1.0] | postpoliomyelitis syndrome (1) [1.0] | parkinsonism (2) [0.1] |
| spaCy | jaundice (1) [1.0] | sideroblastic anemia (2) [0.14] | cascot confidence score measurement (3) [0.11] | susac syndrome (87) [0.1] | parkinsonism (10) [0.1] |
| GUSE | obstructive jaundice (2) [0.09] | blackfan-diamond anemia (2) [0.04] | intelligence (1) [1.0] | mohr-tranebjaerg syndrome (>100) [0.12] | secondary polycythemia (18) [0.19] |
| BlueBERT | jaundice (1) [1.0] | anemia (1) [1.0] | disposition index measurement (>100) [0.08] | tenorio syndrome (6) [0.1] | secondary parkinson disease (1) [1.0] |
| BioSentVec | jaundice (1) [1.0] | anemia (1) [1.0] | intelligence (1) [1.0] | postpoliomyelitis syndrome (1) [1.0] | parkinsonism (3) [0.1] |
| BioBERT | obstructive jaundice (2) [0.09] | anemia (1) [1.0] | cognitive function measureme | postpoliomyelitis syndrome (1) [1.0] | secondary parkinson |

| | | | | | |
|---|---|---|---|---|---|
| | | | nt (>100) [0.11] | | disease (1) [1.0] |
| BLUEBERT-EFO | obstructive jaundice (>100) [0.09] | anemia (disease) (5) [0.15] | mental or behavioural disorder biomarker (>100) [0.11] | syndromic disease (>100) [0.1] | secondary parkinson disease (1) [1.0] |

### Supplementary: Code

**Code Block S1:** Creating an nxontology instance using EFO data

```
from nxontology import NXOntology
def create_efo_nxo(df,child_col,parent_col) -> NXOntology:
    nxo = NXOntology()
    edges = []
    for i,row in df.iterrows():
        child = row[child_col]
        parent = row[parent_col]
        edges.append((parent,child))
    nxo.graph.add_edges_from(edges)
    return nxo
efo_df = pd.read_csv('data/efo_edges_2021_02_01.csv')
efo_nx =
create_efo_nxo(df=efo_df,child_col='efo.id',parent_col='parent_efo.id')
```

**Code Block S2**: Querying the Zooma API

```
zooma_api = 'https://www.ebi.ac.uk/spot/zooma/v2/api/services/annotate'
    payload = {
        'propertyValue':text,
        'filter':'required:[none],ontologies:[efo]'
    }
```

**Code Block S3**: BioSentVec use case example

```
import sent2vec
sent2vec_model = sent2vec.Sent2vecModel()
sent2vec_model.load_model('BioSentVec_PubMed_MIMICIII-bigram_d700.bin')
sentence = 'Psoriatic and enteropathic arthropathies'
sentence_vector = model.embed_sentence(sentence)
```

**Code Block S4**: Google Universal Sentence Encoder v4 use case example

```
import tensorflow_hub as hub
embed = hub.load("https://tfhub.dev/google/universal-sentence-encoder/4")
embeddings = embed([
    "The quick brown fox jumps over the lazy dog.",
    "I am a sentence for which I would like to get its embedding"])
print(embeddings)
```

**Code Block S5**: spaCy and ScispaCy use case example

```
import spacy
nlp = spacy.load("en_core_web_lg")  # make sure to use larger package!
doc1 = nlp("I like salty fries and hamburgers.")
doc2 = nlp("Fast food tastes very good.")
```

```
# Similarity of two documents
print(doc1, "<->", doc2, doc1.similarity(doc2))
# Similarity of tokens and spans
french_fries = doc1[2:4]
burgers = doc1[5]
print(french_fries, "<->", burgers, french_fries.similarity(burgers))
```

**Code Block S6**: BLUBERT and BioBERT use case example

```
from bert_serving.client import BertClient
ip = "localhost"  # subject to how the model API is set up
port = "8888"
port_out = "8889"
text = "body mass index"
with BertClient(ip=ip, port=port, port_out=port_out, output_fmt="list") as
client:
    embeddings = client.encode(text)
    print(embeddings)
```

**Code Block S7**: BLUEBERT-EFO use case example

```
import torch
from transformers import AutoConfig, AutoModelForSequenceClassification,
AutoTokenizer

BASE_MODEL_NAME = "bionlp/bluebert_pubmed_mimic_uncased_L-12_H-768_A-12"
# e.g. "bluebert-efo/pytorch_model.bin"
MODEL_BIN_PATH = "path/to/model.bin"
MODEL_CONFIG_PATH = "path/to/config.json"

config = AutoConfig.from_pretrained(MODEL_CONFIG_PATH, num_labels=1)
model = AutoModelForSequenceClassification.from_pretrained(MODEL_BIN_PATH,
config=config)
tokenizer = AutoTokenizer.from_pretrained(MODEL_BIN_PATH)

text_list_1 = ["body mass index"]
text_list_2 = ["coronary heart disease"]

encodings = tokenizer(text_list_1, text_list_2)
with torch.no_grad():
    scores = model(**encodings)["logits"].reshape(-1).tolist()
    print(scores)
```

### Supplementary: Files

| ID | Name | Description | Count |
|---|---|---|---|
| **S1** | s1_efo_nodes_2021_02_01.csv | EFO node data | 25,390 |
| **S2** | s2_efo_edges_2021_02_01.csv | EFO edge data | 43,132 |
| **S3** | s3_ebi-ukb-cleaned.csv | Cleaned UK Biobank to EFO data | 1,191 |
| **S4** | s4_ebi_exact.tsv | Cleaned UK Biobank to EFO data (Exact) | 530 |
| **S5** | s5_manual_sample.csv | Manual selected UK Biobank traits | 43 |
| **S6** | s6_all-spread.csv | Top 20 EFO mappings with highest EFO-Batet standard deviation across models | 20 |
| **S7** | s7_all-low.csv | EFO mappings where no model had EFO-Batet score >0.95 | 79 |
| **S8** | s8_all-high.csv | EFO mappings where all models had EFO-Batet scire >0.95 | 64 |
